# Actomyosin-driven tension at compartmental boundaries orients cell division independently of cell geometry *in vivo*

**DOI:** 10.1101/397893

**Authors:** Elena Scarpa, Cédric Finet, Guy Blanchard, Bénédicte Sanson

## Abstract

During animal development, planar polarization of the actomyosin cytoskeleton underlies key morphogenetic events such as axis extension and boundary formation. Actomyosin is enriched along compartment boundaries during segmentation of the *Drosophila* embryo, forming supracellular contractile cables that keep cells segregated at boundaries. Here, we show that these contractile actomyosin cables bias the orientation of division in cells in contact with compartment boundaries. By decreasing actomyosin cable tension locally using laser ablation or, conversely ectopically increasing tension using laser wounding, we demonstrate that localised subcellular force is necessary and sufficient to orient mitoses *in vivo.* Moreover this bias is independent of cell geometry and involves capture of the spindle pole by the actomyosin cortex.

## Introduction

Regulation of the orientation of cell division is important for tissue morphogenesis, and if defective, can lead to disease such as tumorigenesis ^1^, kidney malformations ^2^ or microcephaly ^3^. In developing epithelia, mitoses are usually oriented in the plane of the tissue, contributing to tissue elongation ^4^ or homeostasis ^5, 6^. More than 100 years ago, O. Hertwig observed that animal cells often orient their division plane perpendicular to the longest axis of interphase cell shape ^7, 8^. Recently, the distribution of the tricellular vertexes in the fly notum epithelium was found to be also a predictor of the orientation of the division plane and, in moderately elongated cells, a better predictor than interphase cell shape ^9^.

Recent work on isolated cells cultured on micropatterns has shown that physical forces may control the orientation of the mitotic spindle independently of cell shape ^10^. *In vivo*, it has been observed that tissue-level extrinsic forces can orient cell divisions perpendicular to the direction of the stress ^5, 6, 10, 11^. However, because tissue-scale forces also cause planar cell elongation in epithelia ^6, 11-13^, it has been difficult *in vivo* to disentangle a direct effect of forces from an indirect effect on cell geometry.^11^.

Here, we have discovered a population of cells in the *Drosophila* embryonic epidermis whose mitoses do not follow the long axis rule. These cells are located at the parasegmental boundaries and divide perpendicular to a contractile actomyosin cable which forms at the boundary cell-cell interfaces ^14^. We provide evidence that the orientation of the division plane of the boundary cells is governed directly by local tension anisotropy and not by cell geometry or genetic cues.

## Results

### Cells dividing at parasegment boundaries do not follow the long axis rule

During *Drosophila* embryogenesis, the epidermis undergoes waves of cell divisions at extended germ-band stages 9 to 11 ^15,16^. Parasegmental boundaries (PSBs) form through patterning mechanisms and prevent cells or their descendants from changing compartments ^14, 17^ (Fig. 1a). Here we find that at these stages, cells contacting the boundaries (boundary cells, BC) bias their orientation of division differently from non-boundary cells (NBC) (Fig. 1a,b,c). In fixed embryos, NBCs divide predominantly perpendicular to the antero-posterior (AP) axis of the embryo (Fig. 1b,d. Supplementary Fig. 1a,b) In contrast, BCs orient their divisions parallel to the AP axis of the embryo, perpendicular to the PSBs (Fig. 1c,e). This was equally true for cells on both sides of boundary (ie. both *wingless* and *engrailed*-expressing) (Supplementary Fig. 1c,d).

**Figure 1.**
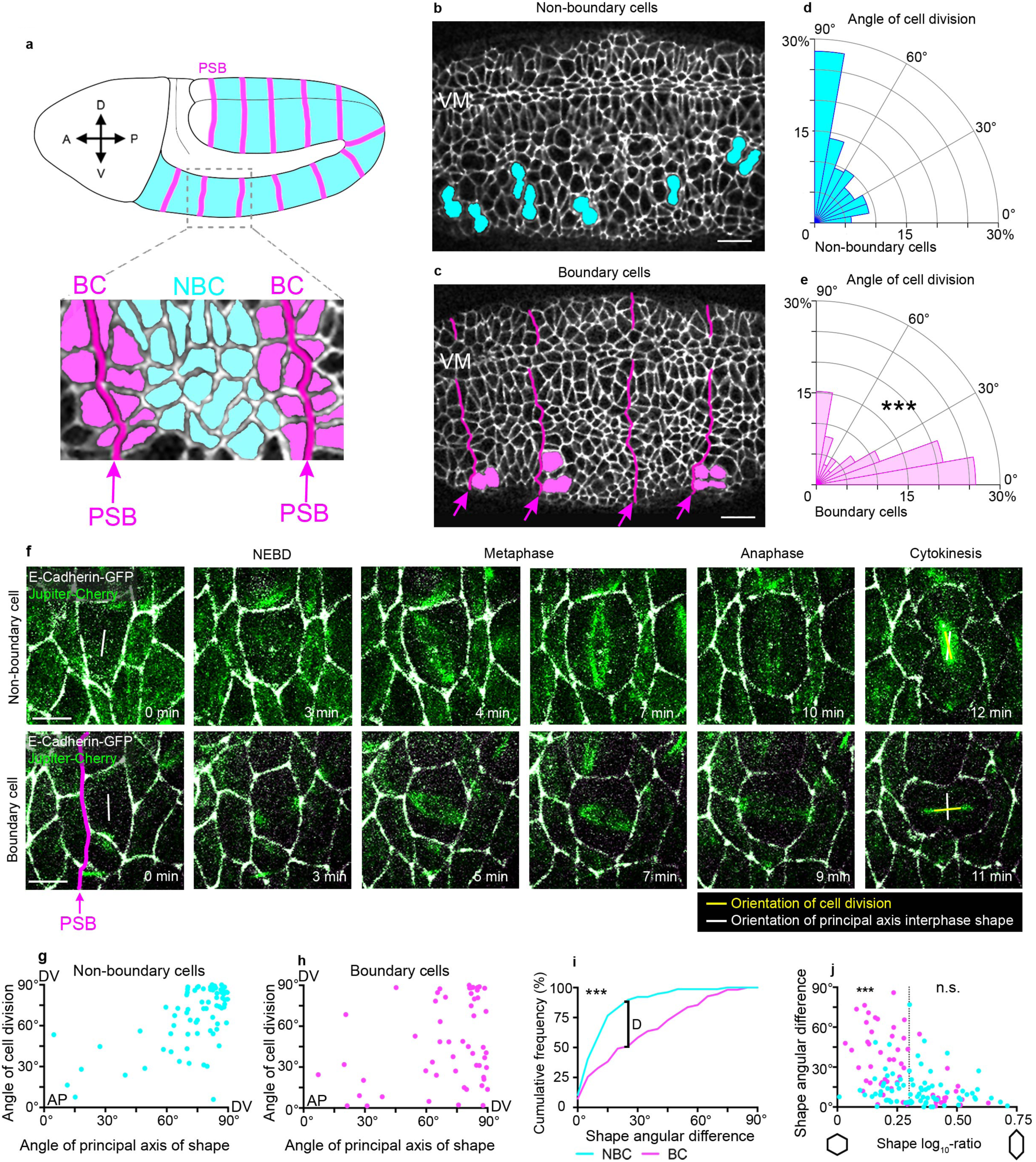
Cells dividing at the parasegment boundary do not follow the long axis rule. **a,** Diagram of a *Drosophila* embryo when the germ-band (blue) is extended (stages 9 to 11). Cell divisions occur throughout the extended germ-band epidermis. The metameric subdivisions are the parasegments, separated by parasegment boundaries (PSBs, pink). BC, boundary cells; NCB, non-boundary cells. **b,c,** Examples of the planar cell division biases found in non-boundary and boundary cells. VM, ventral midline. Scale bar=10 µm. **d,e,** Quantification of the angle of cell division in fixed embryos relative to the antero-posterior axis in both populations (NBC, *n*=391 cell divisions; BC, *n*=289; Mann-Whitney test, *U*=34501, P<0.0001). **f,** Examples of dividing NBC and BC from time-lapse images of a *ubi-DE-Cadherin/en-Venus; jupiter::mCherry/+* embryo. en-Venus was used to identify PSBs (not shown) and jupiter::mCherry (green) to label the mitotic spindle. The orientation of cell division (yellow vector) versus the orientation of interphase shape (white vector) is shown. Scale bar 5 µm. **g**, In NBC, there is a correlation between these two angles, suggesting that these cells follow the long axis rule (*n*=77; Spearman’s rho test, *r*=0.48, P<0.001). **h**, This correlation is absent for BC (*n*=55; Spearman’s rho test, *r*=0.17, P=0.19). **i,** Cumulative histogram for the angular difference between the orientation of cell division and the orientation of interphase cell shape for NBC (blue) and BC (magenta) (NBC, *n*=77; BC, *n*=55; Kolmogorov-Smirnov test, *D*=0.41, P<0.001). **j**, Angular difference between division and shape as a function of the cell shape log_10_-ratio. Above a log_10_-ratio threshold of 0.3 (DV/AP aspect ratio of 2), both NBC and BC behave similarly (NBC, *n*=48; BC, *n*=16; Kruskal-Wallis test, *H*=39.58, P>0.99), following the long axis rule. Below 0.3 however, NBC and BC behaviours are significantly different (NBC, *n*=29; BC, *n*=39; Kruskal-Wallis test, *H*=39.58, P=0.0003).

Hertwig’s rule says that the plane of cell division is perpendicular to the long axis of the interphase cell ^7, 8, 18^. To ask if cells were following the long axis rule, we imaged dividing cells in a live embryo, using E-Cadherin-GFP to monitor cell shapes, and the microtubule binding protein Jupiter-Cherry^19^ to visualise the mitotic spindle (Fig. 1f). We find that the metaphase spindle rotates within the tissue plane, stabilising at anaphase onset (see Fig. 5a-c). For each cell division, we determined the principal axis of cell shape prior to nuclear envelope breakdown (NEBD) and compared its orientation with the mitotic spindle orientation at anaphase (Fig. 1f-j). We find that 80% of interphase cells have their long axis oriented parallel to the dorso-ventral embryonic axis and this is true of both BCs and NBCs (Supplementary Fig. 1f). However, while NBCs follow the long axis rule, with a strong correlation between angle of cell division and angle of cell shape (*r*=0.48; Fig. 1g), BCs do not (*r*=0.17; Fig. 1h)(Fig. 1i). This might be partly explained by the finding that BCs are less strongly elongated than NBCs (Supplementary Fig. 1g). However, we found that while highly elongated BC divide according to their shapes, the moderately elongated BCs do not follow the long axis rule, in contrast with the NBCs of equivalent aspect ratio (Fig. 1j; Supplementary Fig. 1e,h). These results suggest additional cue(s) other than cell shape control the orientation of BC divisions.

### Actomyosin cables are necessary and sufficient to orient boundary cell divisions

PSB cell-cell interfaces are enriched in actomyosin, and form tissue-scale contractile cables (Fig. 2a,b) that act as mechanical barriers limiting cell mixing caused by cell divisions and, earlier in development, polarised cell intercalations ^14, 20^. Since force anisotropies have been reported to control cell division orientation *in vitro*^10^ as well as in tissues ^5, 6, 11^, we hypothesised that the actomyosin cable at PSBs might act as a source of anisotropic tension during mitosis. As previously reported, live imaging using GFP-tagged Myosin II Regulatory Light Chain (MRLC-GFP) shows that the actomyosin cable-like enrichment persists at the cortex of boundary cells during division (Fig. 2c,d)^14^. We asked whether the actomyosin cable is required for the division orientation bias we observed in these cells. We examined *wingless (wg)* null mutant embryos, where actomyosin fails to accumulate at PSBs ^14, 20, 21^ (Fig. 2e). Strikingly, the majority of BCs now divide along the DV axis like NBCs (Fig. 2e; Supplementary Fig. 2a,b). To test if this difference was caused by the loss of actomyosin enrichment in *wg* mutants, we inhibited Myosin II activity in two different ways. First, we injected wild type embryos with a concentration of the Rok inhibitor Y-27632 that does not affect cytokinesis but does disrupt boundary function ^14, 21^. Second, we overexpressed a dominant-negative form of the Myosin II Heavy Chain in the epidermis ^14, 22^. Both experiments disrupt the division orientation bias in BCs as in *wg* mutants (Fig. 2f; Supplementary Fig. 2c-f). These experiments indicate that the actomyosin cable at PSBs is required for orienting the BCs divisions perpendicular to the boundary. We next asked whether BCs follow the long axis rule when actomyosin contractility is inhibited. We live imaged embryos injected with Y-27632 and examined cell shape orientation prior to division as before. This analysis shows that, indeed, BCs follow the long axis rule in Y-27632 but not H_2_O injected embryos (Supplementary Fig. 2h-j). We further checked that Y-27632 treatment did not affect cell shape orientation or elongation (Supplementary Fig. 2k,l). These results indicate that in absence of a contractile actomyosin cable at the boundary, BCs behave as NBCs.

**Figure 2.**
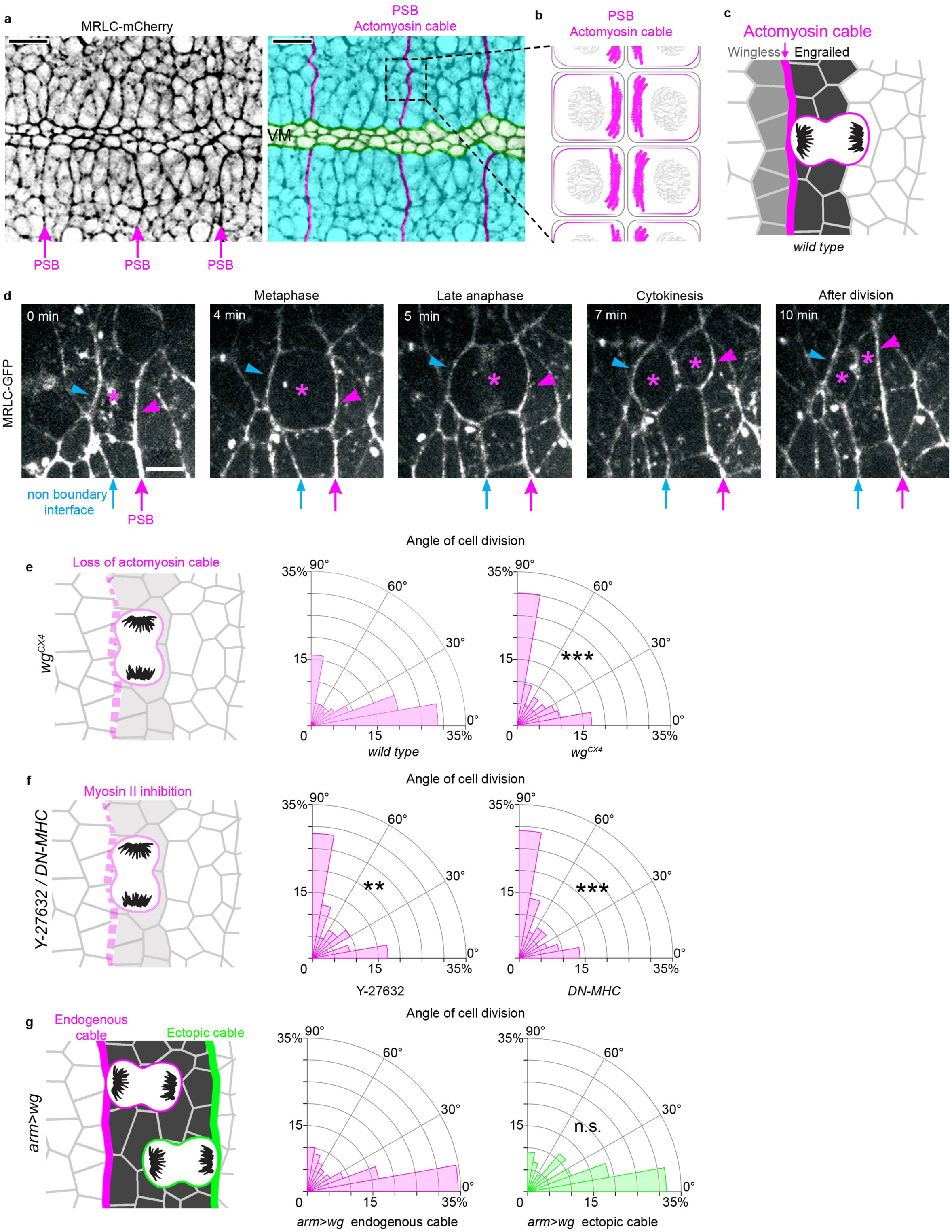
The actomyosin cable at the PSB is necessary and sufficient to orient boundary cell division. **a,** Image from a MRLC-Cherry movie. Arrows label actomyosin cables at PSBs. Colour-coded version of the same image: PSB actomyosin cables are highlighted in magenta, extended germband tissue in cyan, ventral midline (VM) in green. **b**, Diagram representing a zoomed-in PSB actomyosin cable. The cable is formed by apposed actomyosin-enriched cell cortices on either side ^14^. **c,** Diagram representing AP-oriented cell divisions at the PSB. **d,** The actomyosin cable at the PSB is maintained throughout cell division. Time-lapse images of a dividing cell in a MRLC-GFP expressing embryo. Asterisk indicates the dividing cell and its daughters. Magenta arrow: PSB, blue arrow: non-boundary interface. Scale bar 5 µm. **e,** The actomyosin cable is required for OCD in boundary cells. Radial histograms of OCD in WT embryos or *wg^CX4^* mutants (both quantifications for stage 9 to 11 embryos) (WT, *n*=418; *wg^CX4^, n*=378; Mann-Whitney test, *U*=58262, P<0.0001). **f,** Radial histograms of OCD in embryos injected with the Rok inhibitor Y-27632 (*n*=81; Mann-Whitney test, *U*=2548, P=0.0073) or expressing DN-MHC (*n*=238; Mann-Whitney test, *U*=23586, P<0.0001). **g,** Cell divisions orient perpendicular to the AP axis when an ectopic actomyosin cable is generated by ubiquitous Wg expression. Compare histograms of OCD at endogenous PSB (left, *n*=218) with OCD of cells in contact with the ectopic cable (right, *n*=133; Mann-Whitney test, *U*=14489, P=0.994).

To ask then if a cable-like actomyosin enrichment is sufficient to orient cell division, we generated an ectopic cable at the posterior interface of the Engrailed expression domain by uniformly expressing Wg in the epidermis ^21^ (Fig. 2g). We have shown previously that this interface behaves as an ectopic PSB, with a similar actomyosin enrichment and increased interfacial tension as for endogenous PSBs ^21^. We find that cells contacting this ectopic cable orient their division perpendicular to it, as at the endogenous cable (Fig. 2g; Supplementary Fig. 2g). Taken together, the above findings indicate that a contractile actomyosin cable is both necessary and sufficient to orient boundary cell division.

### Elevated tension at PSB actomyosin cables is required for orienting cell division

Since PSB actomyosin cables act as mechanical barriers preventing cell mixing during body axis elongation ^20^ and segmentation ^14^, we postulated they might orient cell divisions as a consequence of their mechanical properties. Cortical tension can be estimated by severing cell-cell junctions and comparing recoil velocities, assuming that friction is the same for all junctions ^23, 24^. Using this approach, we have shown previously and confirm here that cortical tension at PSB junctional interfaces is elevated, about two-fold that of non-PSB interfaces (Supplementary Fig. 3a-f) ^20, 21^. Based on these measurements, we hypothesised that cells at the boundary might experience a cortical anisotropy in tension that biases their division orientation^10^.

We developed a strategy to decrease tension locally at the PSB, in order to ask if this would change the division orientation of boundary cells in contact with the cable. We reasoned that a single cut in the actomyosin cable could decrease tension along some of its length. To check if this assumption was correct, we ablated the cable twice, leaving an interval of 20 seconds before the second cut was performed one cell diameter away further along the cable (Fig. 3a-b)^25^. Comparison of recoil velocities at first and second ablation sites show a nearly two-fold decrease in recoil speed, indicating that a single cut effectively reduces tension in the cable (Fig. 3g-i). This is consistent with contractile stress being propagated and integrated along actomyosin-enriched supracellular cables^26^. Based on this finding, we cut the PSB actomyosin cable in proximity of boundary cells at metaphase, when the spindle still rotates (see Fig. 5a-c), to ask if a tensile cable was required for their division orientation. To efficiently ablate the PSB cable for the duration of the division, we repeated the laser ablation every 25 seconds to suppress actomyosin-mediated wound healing ^27^. Kymographs were inspected to check that no repair occurred and that the cable had lost tension throughout the experiment (Supplementary Fig. 3j). As a control, we targeted the cable near metaphase cells using low intensity light (Fig. 3d). We show that loss of tension at the actomyosin cable significantly reduced the ability of BCs to orient their divisions perpendicular to the cable, compared to control cells (Fig. 3d-i), despite their cell shapes being indistinguishable from those of control BCs (Supplementary Fig. 3k,l). Note that upon ablation, we find only a marginal decrease in Myosin II along the cable (about 10%, Supplementary Fig. 3i, see also Discussion), suggesting that it is the loss of tension, rather than a loss of actomyosin enrichment, which is the important cue. In summary, these experiments indicate that a cortical anisotropy of tension at the PSB is necessary to orient cell divisions *in vivo*, independently of cell shape.

**Figure 3.**
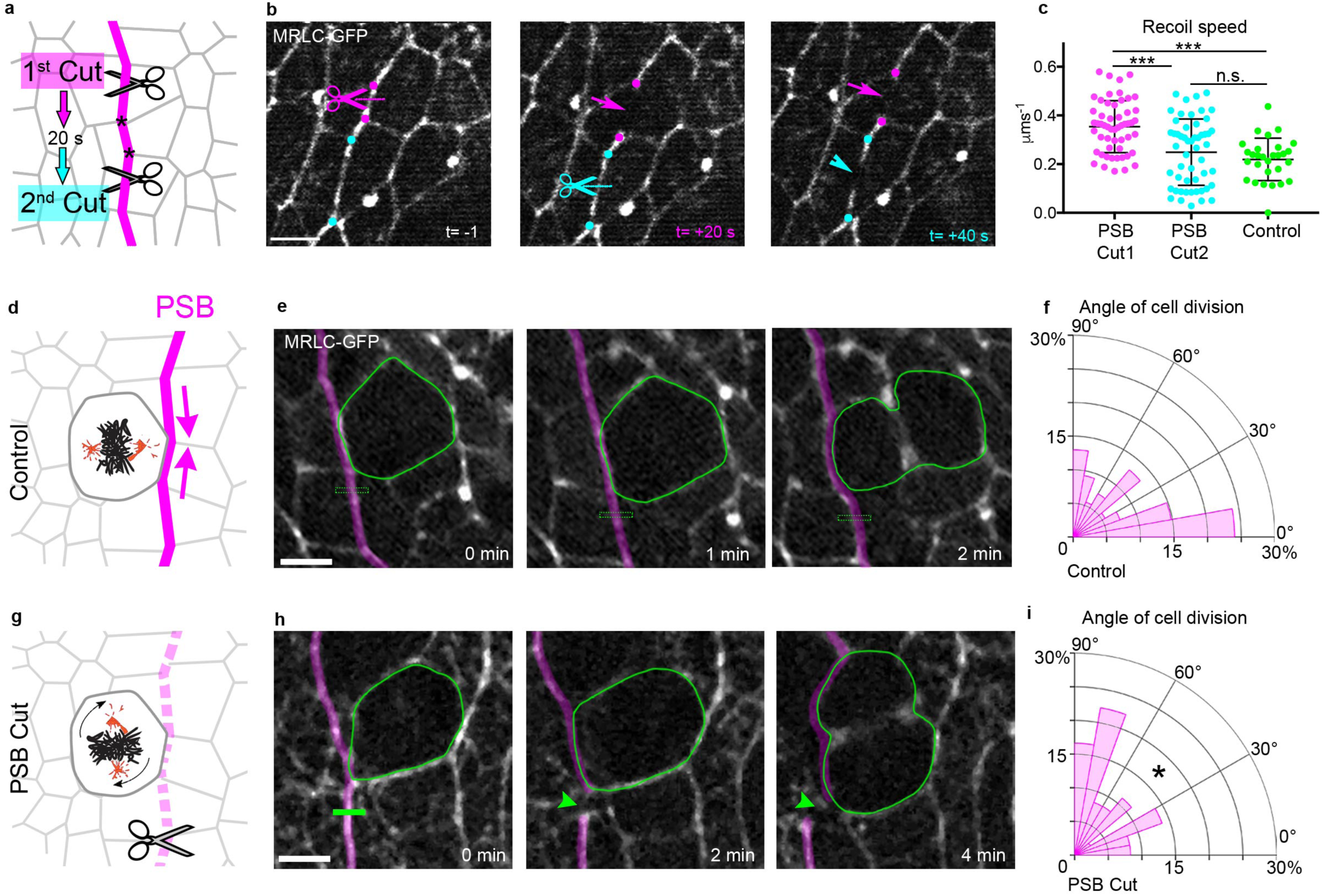
Anisotropic local tension at the actomyosin cable is required for cell division orientation of boundary cells. **a-c** Tension is non-autonomously integrated over adjacent interfaces at the PSB actomyosin cable. (a) The actomyosin cable is cut once and allowed to relax. After 20s, when relaxation is maximal, a second cut is performed at a distance of two junctions (asterisks). (b) Still images from a consecutive ablation experiment. The site of the first cut and its recoiling cut ends are highlighted in magenta, while the second cut site is highlighted in cyan. Scale bar 5 µm. (c) Speed of recoil upon ablation (PSB Cut1, *n*=52; PSB Cut2, *n*=52; Control, *n*=29; Kruskal-Wallis tests; PSB Cut1 vs PSB Cut2, *H*=26.44, P=0.0002; PSB Cut1 vs Control, *H*=26.44, P<0.0001; PSB Cut2 vs Control, *H*=26.44, P=0.573). Means ± SDs shown. **d,g** To impair local tension at the PSB, the actomyosin cable is cut next to a mitotic boundary cell in metaphase and the orientation of the cell division is measured. Actomyosin cables were either control treated with the same (25%) irradiation used for image acquisition (d) or cut by 100% 927 nm laser (g). **e,h** Still images from a loss of PSB tension experiment. PSB actomyosin cable is highlighted in magenta. Green rectangle, site of control treatment or of ablation. Green arrowhead highlights cable recoil. Scale bar 5 µm. **f,i** Histograms displaying cell division orientation for Control and PSB-ablated cells (Control, *n*=54; PSB cuts, *n*=36; Mann-Whitney test, *U*=717, P=0.0356).

### Boundary cells do not require Pins, Mud or the vertex rule to orient their divisions

In mammalian cells, E-Cadherin recruits the cortical regulator LGN (Pins in *Drosophila*)^28^ to adherens junctions ^29^. When epithelial monolayers are stretched, LGN/Pins is used to align cell division orientation with the force direction, independently of cell shape ^30^. We therefore investigated Pins requirement for cell division orientation at PSBs. Knockdown of Pins by RNAi (Supplementary Fig. 4d-f) did not significantly change cell division orientation for either BCs or NBCs (Supplementary Fig. 4h,i), despite Pins being required for spindle orientation along the apico-basal axis of the epithelium (Supplementary Fig. 4g)^28^. These results suggest that Pins is not required to orient BC divisions at the PSB or even in NBC divisions away from the PSB.

In the *Drosophila* notum, cells align their divisions according to the spatial distribution of their tricellular vertices ^9^. Forces influence these vertex positions, and in turn vertices orient mitoses by anchoring astral microtubules to the cell cortex via Mud, the NuMA homologue ^9^. In this tissue, vertex distribution is a better predictor of the orientation of division than cell elongation^9^. PSBs are straighter than non-boundary columns of cell contacts ^14, 20^ and this straightness is predicted to promote clustering of vertices along the PSB (Fig. 4a,a’), so it seemed plausible that vertex distribution might explain the orientation of BC divisions. To test this, we imaged live embryos expressing E-Cadherin-GFP ^31^ and MRLC-mCherry ^32^ and tracked 160 dividing cells (See Methods). Although the tracked BCs do have a distribution of vertices biased by the boundary as we predicted (red box, Fig. 4b), the distribution of vertices does not do better than the long axis rule at predicting cell division orientation in those cells (Fig. 4c; Supplementary Fig. 4j-l).

**Figure 4.**
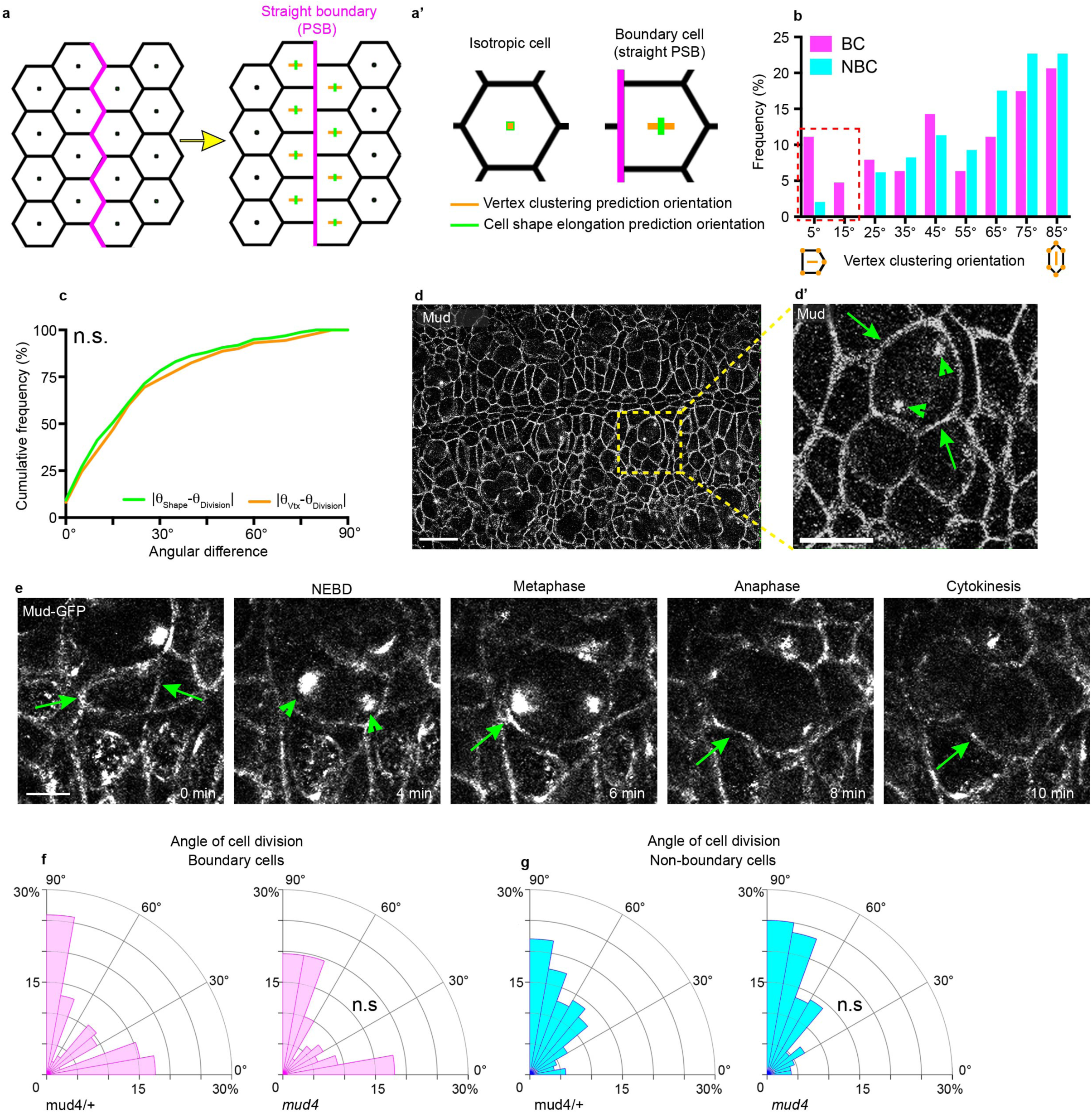
Boundary cells do not require the vertex rule to orient their divisions. **a-a’,** (a)For a given hexagonal starting configuration of cells (left panel), imposing a straight boundary (PSB, thick magenta line, right panel) introduces both clustering of tricellular vertices perpendicular to the boundary and a small cell shape elongation parallel to it. (a’, inset of a), While for an isotropic hexagonal cell the vectors η_Vertex_ and η_Shape_ equal 0, for cells with a straight boundary η_Vertex_ and η_Shape_ diverge in their orientation and magnitude, with η_Vertex_ perpendicular to the boundary and η_Shape_ parallel to it (vertex clustering value, η_Vertex_=0.21; shape elongation value, η_Shape_=0.13). Thus a straight multicellular interface could in itself introduce a cell division orientation bias if the orientation mechanism depended on the clustering of cell vertices. **b,** Histogram of vertex cluster orientation in relation to the anteroposterior axis of the embryo for BC and NBC at t=-12 mins from cytokinesis from E-Cadherin-GFP,MRLC-mCherry movies (n=97, NBC; n=63, NBC; Mann-Whitney test, *U*=2593 P=0.107). **c**, Cumulative histogram of the shape and vertex angular differences for all cell divisions in neurectoderm (*n*=160 from 3 embryos; Kolmogorov-Smirnov test, *D*=0.075, P=0.759). **d-d’**, Immunostaining of a stage 9 embryo with anti-Mud antibody. Scale bar 20 µm. d’ inset from d. Arrowheads, endogenous Mud localisation at mitotic spindle poles; arrows, localization of Mud at bi-cellular junctions. Scale bar 10 µm. **e,** Stills of a dividing cell from a mud-GFP expressing embryo. Arrowheads, endogenous Mud localisation at mitotic spindle poles; arrows, localization of Mud at bi-cellular junctions. Scale bar 5 µm. **f,** Histogram of cell division orientation for boundary cells (BC) in *mud^4^* (*n*=85), or *mud^4^/+* (*n*=66; Mann-Whitney test, *U*=2786, P=0.943) embryos. **g,** Histogram of cell division orientation for non-boundary cells in *mud^4^* (*n*=128), or *mud^4^/+* (*n*=172; Mann-Whitney test, *U*=9811, P=0.107) embryos.

In line with the above result, we find that Mud is not localised at vertices in the embryo (Fig. 4d-e), but in contrast to Pins, Mud is mildly enriched at PSBs (Supplementary Fig. 4a-c), so we tested the requirement of Mud by analysing *mud* null mutant embryos ^33^, in which endogenous Mud is not detected (Supplementary Fig. 4m). Mud controls epithelial cell division orientation along the apico-basal axis ^9, 28^, and indeed we find that 45% of cell divisions occurred out of plane (Supplementary Fig. 4n). Therefore, we restricted our analysis to the remaining 55% of planar mitoses. We find that Mud loss of function did not change cell division orientation for either BCs or NBCs (Fig. 4f-g). Together, these results show that the planar division orientation of boundary cells is neither dependent on Pins/LGN nor does it rely on tricellular vertex distribution of Mud/NuMA.

### PSB actomyosin enrichment restricts spindle pole motility

An alternative explanation for the oriented division in BCs is that Myosin-mediated contractility at the PSB directly impacts spindle pole positioning, especially as Myosin-II is required for spindle pole separation after NEBD, both in mammalian cells ^34^ and in *C. elegans* embryos ^35^. To explore this, we characterized the dynamics of spindle pole motility in BCs versus NBCs. Imaging of mitotic spindles using Jupiter-Cherry revealed a rotational behaviour from NEBD throughout metaphase in both NBCs and BCs, which decreases during the 2 minutes before anaphase onset, when the spindle is stabilised (Fig. 5a-c). As in vertebrate embryos ^36^, the orientation of the mitotic spindle poorly correlated with cell shape at NEBD and this correlation significantly improved at anaphase, indicating that mitotic spindles adopt their final orientation just before anaphase (Fig. 5b). We did not, however, observe any differences in spindle dynamics between BCs and NBCs (Fig. 5c).

**Figure 5.**
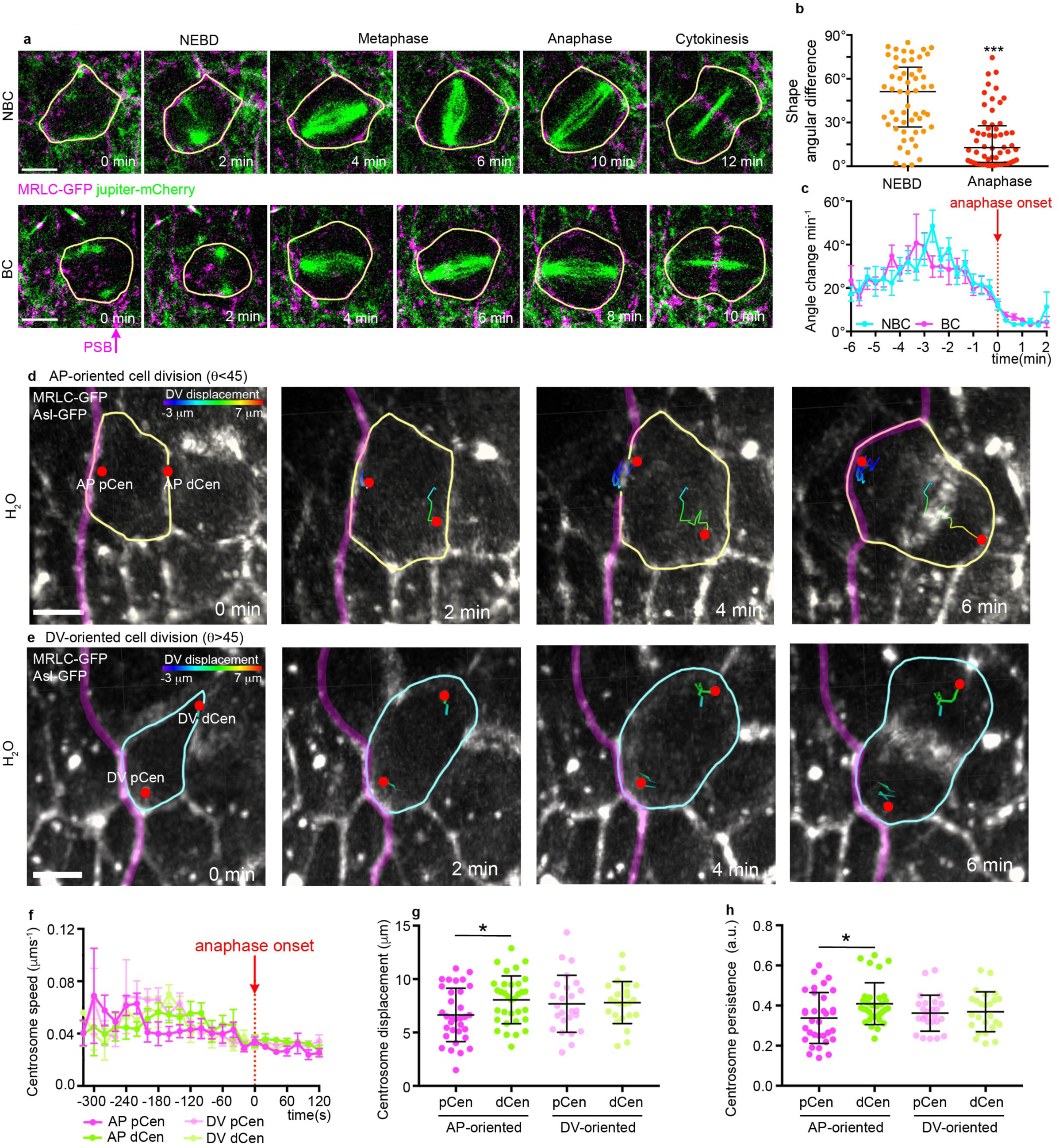
Spindle pole motility is restricted adjacent to the PSB. **a-c** The mitotic spindle of ventral neurectoderm cells is highly dynamic in metaphase, and its orientation stabilises at anaphase onset. (a) Still images from a time-lapse movie of a MRLC-GFP and Jupiter-Cherry-expressing embryo. A Representative non-boundary cell (top) and a boundary cell are shown. Scale bar 5 µm. (b) Shape angular difference at NEBD and anaphase (*n*=56; Kolmogorov-Smirnov test, *D*=0.532, P<0.001). Median±interquartile range shown. (c) Absolute angular rotation per minute of the mitotic spindle for BC (*n*=27) and NBC (*n*=28) from NEBD to cytokinesis. t=0, anaphase onset. Mean ± s.e.m.s shown. **d,e** Representative AP-oriented (d, θ<45) and DV-oriented (e, θ>45) BC cell division from an H_2_O injected embryo expressing MRLC-GFP and Asl-GFP. Centrosome tracks are highlighted and colour-coded for DV displacement. Scale bar 5 µm. **f** Centrosome speed from NEBD to cytokinesis (AP pCen, *n=*16; AP dCen, *n=*16; DV pCen, *n=*19; DV dCen, *n*=19). t=0, anaphase onset. Mean ± s.e.m. shown. **g** Total displacement for each centrosome from NEBD to cytokinesis (AP pCen, *n=*33; AP dCen, *n=*33; DV pCen, *n*=25; DV dCen, *n*=25; One-way Anova, *F*=2.252, P=0.059; AP pCen vs AP dCen, P=0.0317; all other comparisons P>0.30). Means ± SDs shown. **h** Persistence for each centrosome from NEBD to cytokinesis (AP pCen, *n=*33; AP dCen, *n=*33; DV pCen, *n*=25; DV dCen, *n*=25; One-way Anova, *F*=2.863, P=0.040; AP pCen vs AP dCen, P=0.0241; all other comparisons P>0.20). Means ± SDs shown.

Next, we asked whether the mitotic spindle pole closest to the PSB behaved differently from the pole more distal from the PSB. To separately track spindle poles, we imaged embryos expressing the centrosome marker Asl-GFP ^37^ and MRLC-GFP to identify the PSB. Tracking spindle poles from NEBD to cytokinesis (Fig. 5d,e) revealed that the speed and the distance covered by poles throughout mitosis is comparable (Fig. 5f, Supplementary Fig. 5a). However, in BCs dividing perpendicular to the boundary (AP-oriented, Fig. 5d), the centrosome proximal to the PSB (pCen) was displaced less (Fig. 5g) and displayed a trajectory significantly less persistent (Fig. 5h) than the centrosome distal from the PSB (dCen). In contrast, proximal and distal centrosomes of BCs dividing without a bias toward the boundary (DV-oriented, Fig. 5e), behave similarly (Fig. 5g-h; Supplementary Fig. 5b,c). Moreover, in Y-27632 treated embryos, each centrosome displays similar motility even in BCs that divide perpendicular to the PSB (Supplementary Fig. 5d-i). This suggests that the PSB, through the action of Myosin II, orientates BC cell division by capturing the more proximal centrosome and limiting its motility.

### Ectopic tension anisotropy is sufficient to orient cell divisions

Our results so far indicate that tension anisotropy caused by actomyosin cables can orient cell division via a centrosome capture mechanism. To ask if tension anisotropy is sufficient to orient mitoses *in vivo*, we sought to change tension in the vicinity of NBCs, by laser wounding the nearby epithelium to provoke a wound healing response^5, 27^. We found fast Myosin II accumulation at small wounds, peaking at about 90 seconds after wounding (Fig. 6 a,b). To ask how these changes might be associated with tension, we ablated the repairing region where medial actomyosin accumulates and measured recoil (Fig. 6c). Myosin II enrichment after wound healing in response to ablation recoils significantly faster than in control junctions (Fig. 6d). Thus, using a treatment that does not have any detectable detrimental effects on neighbouring cells (for example, we do not observe any cell delamination, Supplementary Fig. 6a,b), we can locally increase tension with good spatial and temporal resolution.

**Figure 6.**
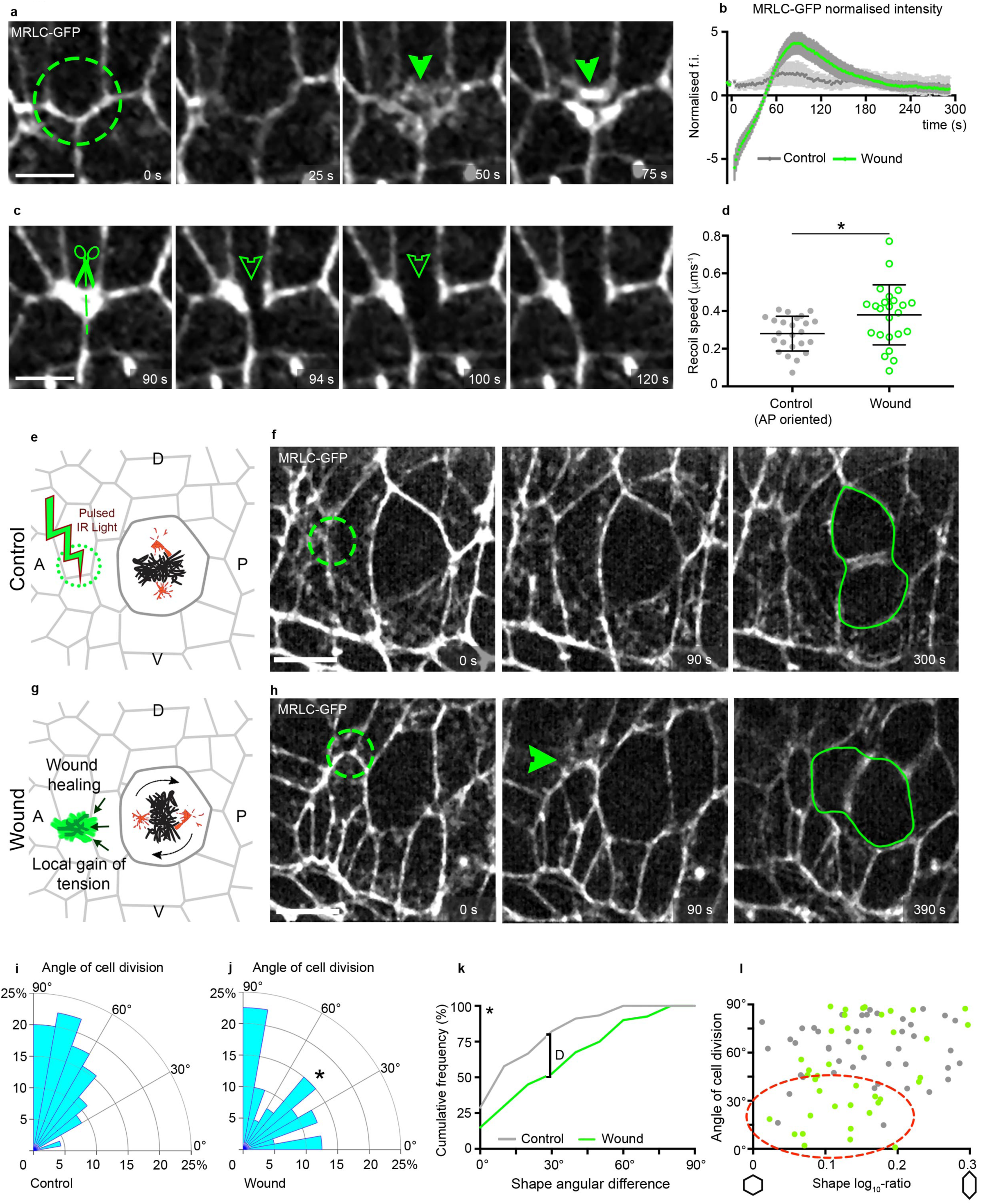
Anisotropic local tension generated by IR laser wounding is sufficient to reorient cell divisions of non-boundary cells. **a,** IR irradiation of adherens junctions results in a wound healing response with transient accumulation of a medial MRLC-GFP meshwork. Scale bar 5 µm. **b** Normalised fluorescence intensity of the irradiated area over time (*n=*19, sham; *n=*26, wound). Myosin intensity peaks at t=90s after irradiation. **c,** Laser ablation of the myosin meshwork displays rapid recoil. Scale bar 5 µm. **d,** Recoil speed upon laser cut of the healing myosin mesh, compared to AP oriented unwounded junctions (*n=*24, Control; *n=*24, Wound; Student’s *t*-test, *t*=2.65, P= 0.016). Means ± SDs shown. **e,g** To increase tension locally, a laser wound is generated by IR irradiation of a circular ROI oriented in AP at one cell distance from a mitotic non-boundary cell in metaphase (e). Wound healing results in generation of a local gain of tension (g). The orientation of the cell division is measured. The tissue was either control treated with the same (25%) irradiation used for image acquisition (e) or irradiated at 80% 927 nm laser power (g). **f,h** Still images from a gain of tension experiment. Green circle, site of wounding. Green arrowhead highlights myosin accumulation upon wounding. Scale bar 10 µm. **i,j** Histograms of OCD for sham and wounded cells (*n*=45, Control; *n*=40, Wound; one-tailed Mann-Whitney test, *U*=682, P=0.027). **k** Cumulative frequency distribution of the difference between the orientation of cell division and the orientation of the principal axis of cell shape (*n*=45, Control; *n*=40, Wound; Kolmogorov-Smirnov test, *D*=0.3278, P=0.0211). **l** Cell division orientation as a function of cell elongation (cell shape log_10_-ratio). Red circle highlights treated AP-oriented cells (θ_midline_ < 45°) that are poorly elongated (cell shape log_10_-ratio < 0.2) (*n*=45, Control; *n*=40, Wound; one-tailed Mann-Whitney test, *U*=682, P=0.027).

With this new tool in hand, we sought to locally increase tension at the vicinity of a metaphase NBC, either on its anterior or posterior side (Fig. 6g-h). Controls were performed by irradiating a region of the same size with low light (Fig. 6e,f). Whereas control cells divide mostly along DV as expected (Fig. 6f,i), we find that, in contrast, NBCs next to an A or P source of tension divide more frequently along the AP axis (Fig. 6h,j), without changing their orientation or elongation (Supplementary Fig. 6c,d). NBCs near a wound also no longer followed the long axis rule as well as control cells did (Fig. 6k). We conclude that local anisotropies in Myosin II-associated tension are sufficient to orient cell division *in vivo*. Furthermore, the proportion of cells orienting relative to the source of ectopic tension is greatest for moderately elongated cells, suggesting a competition between this cue and the long axis cue (red oval, Fig. 6l).

## DISCUSSION

Our study demonstrates that *in vivo*, an anisotropy in actomyosin-mediated tension at the cortex of dividing cells can orient their plane of division independently of cell geometry. This is a novel finding that adds an important insight into *in vivo* studies of cell division where it is unclear if forces act directly, or indirectly via a change in cell geometry, because tissue-scale forces elongate cells ^5, 6, 38^.

We find that in all the experimental conditions where cells have one side of the cortex belonging to a tensile actomyosin-enriched parasegmental boundary, their division orientation are biased by the boundary rather than by the cells’ longest axis. This is true unless the boundary cells are very elongated (Fig. 1j), in which case the mechanical boundary is unable to reorient the dividing cells. We observed the same result in our wounding assay to generate high tension locally: the cells that are not reoriented are the most elongated (Fig. 6l). This suggests that above a certain threshold of cell elongation, which we estimate being around 2 for the ratio between long axis and short axis (Fig. 1j; Supplementary Fig. 1h), the long axis rule ‘wins’ over localised cortical actomyosin tension. This is likely to explain the bimodal distribution of division orientation we observe in our fixed data: although a high proportion of BCs divides perpendicular to the boundary (along AP), there is always a group of cells that still divides along DV (Fig. 1e). Imaging live embryos, we show these indeed are the strongly elongated cells (Supplementary Fig. 1h). We also find systematically that when tension is disrupted at the PSBs, the boundary cells now divide according to the orientation of their long axis, which is on average along DV (Fig. 3i, Supplementary Fig. 3k). This suggests that the default orientation cue in NBC tissue is the cell’s shape. It is still unclear why cells tend to divide according to their long axis. For strongly elongated cells, it is possible these do not manage to round up completely and that steric hindrance of the spindle constrains its orientation ^39, 40^. For moderately elongated cells, such explanation is unlikely. It was recently discovered that the spatial distribution of vertices in the *Drosophila* notum epithelium predicts the orientation of the division better than the long axis rule, in particular for moderately elongated cells ^9^. The explanation for this is that Mud (the homolog of NuMA) interacts with septate junction (similar to tight junctions) components at vertices, causing a redistribution of forces exerted by the astral microtubules. We have ruled out a role for Mud in experiments in this paper, suggesting that the default orientation of cell division for NBC is determined by other mechanisms. Direct force sensing could be involved, although we cannot disentangle this from the long axis rule in NBCs.

In BCs, our results points toward a direct role of actomyosin-mediated tension anisotropy in biasing spindle orientation. We found that the *Drosophila* homolog of LGN, Pins, is not required for the orientation bias in BCs (or NBCs). Consistent with this, Myosin II activity was shown not to affect Pins localization in *Drosophila* embryos ^41^. Since Mud is also not required, this suggests that the Pins/Mud/Dynein complex does not play a role in our example ^18^. It is therefore possible that a mechanosensitive pathway in the cells is what responds to tension and interacts with the astral microtubules to orient the spindle. Such a molecular cue could be E-Cadherin ^42^, whose localisation can change when tension at the cortex is decreased or increased^43^. Illustrating this possibility, E-Cadherin/LGN interactions control cell division orientation in response to tissue stretch in mammalian cells ^30^. Another component of adherens junctions, the protein Canoe/Afadin, is required to attach the actomyosin network to the cortex in early Drosophila embryos ^44^. It promotes spindle orientation by recruiting cortical actin via RhoA and the formin Dia in *Drosophila* sensory organ precursors ^45^. Relevant to this, one of its zebrafish homologs, ZDia2, is required for spindle orientation towards a cortical enrichment of actin during epiboly ^46^. Future work could test if tension-dependent localisation of adherens junctions components such as Canoe or E-Cadherin is important for both force integration and spindle orientation at the PSB.

We have found that the centrosome proximal to the actomyosin cable is significantly less mobile during metaphase, suggesting it might be captured by the cortex experiencing high tension. Supporting this idea, cortical actomyosin contractility has been recently shown to promote clustering of centrosomes by limiting their motility in cultured cancer cells with supernumerary centrosomes ^47^. Forces can capture the mitotic spindle poles via subcortical actin clouds ^10^ or the actin-microtubule-binding motor Myosin10 ^48^. So far, actin clouds have only been reported in cultured cells ^10, 48^ and in *Xenopus* epithelia ^49^ and Myosin10 is not expressed during early *Drosophila* embryogenesis ^50, 51^. Alternatively, actomyosin-mediated tension at the PSBs might cause an anisotropy in cortex stiffness, which could bias the balance of forces orienting the spindle. Indeed, Myosin II, together with the actin-membrane crosslinker Moesin, is essential for cortex stiffening during mitotic cell rounding ^41, 52^. Moreover, impairment of cortical stiffening by actin-depolymerizing drugs or Myosin II inhibition perturbs cell division orientation *in vivo* ^41, 53, 54^. A stiff enough actomyosin cortex is essential to balance the tension that cortical force generators exert when pulling on the spindle in *C. elegans* ^55^ and to drive asymmetric spindle localization in mouse oocytes ^56^.

Our study uncovers a novel effect of actomyosin supracellular cables in orienting cell divisions *in vivo*. Actomyosin cables are very common in developing epithelia and are found not only at compartmental boundaries but also during tissue closure, wound healing, tube formation and convergence extension of tissues ^57^. In the wing disc, where actomyosin supracellular tension has been identified at both AP and DV boundaries, no orientation of cell division perpendicular to the boundary has been reported ^58-60^. However, the tension at the AP boundary is controlled cell-autonomously by Hedgehog signalling ^25^. In the embryo, in contrast, we find that severing the actomyosin cable causes tension to be lost non-cell autonomously along the cable. We also find that Myosin II is decreased along the cable as a consequence of this loss of tension (Supplementary Fig. 3i). This indicates that in the embryo, actomyosin cables are supracellular tensile structures, where Myosin II enrichment might be reinforced by a mechanosensitive feedback ^26^. In contrast with the wing disc, the embryonic ectoderm is a simple epithelium lacking septate junctions or extracellular matrix, where force transmission along an actomyosin cable might be facilitated. This might in turn influence how much tension is generated and how much it can bias division orientation.

What function might tension-generated orientation of cell division serve during morphogenesis? In our example, cell divisions start towards the end of axis extension ^16^, when large-scale polarised cell intercalations have generated a disordered epithelium ^61^. It is possible that this intense period of cell division is required to restore optimal tissue packing ^62, 63^. In this context, the separation of metameric units by mechanical barriers that bias cell division in a tension-dependent manner might be key for the conservation of shape between parasegments. More generally, our study raises the possibility that cell division orientation biases directly caused by local tension anisotropies could be important in the maintenance of correct tissue and organ sizes during growth.

## SUPPLEMENTARY FIGURE LEGENDS

**Supplementary Figure 1:**
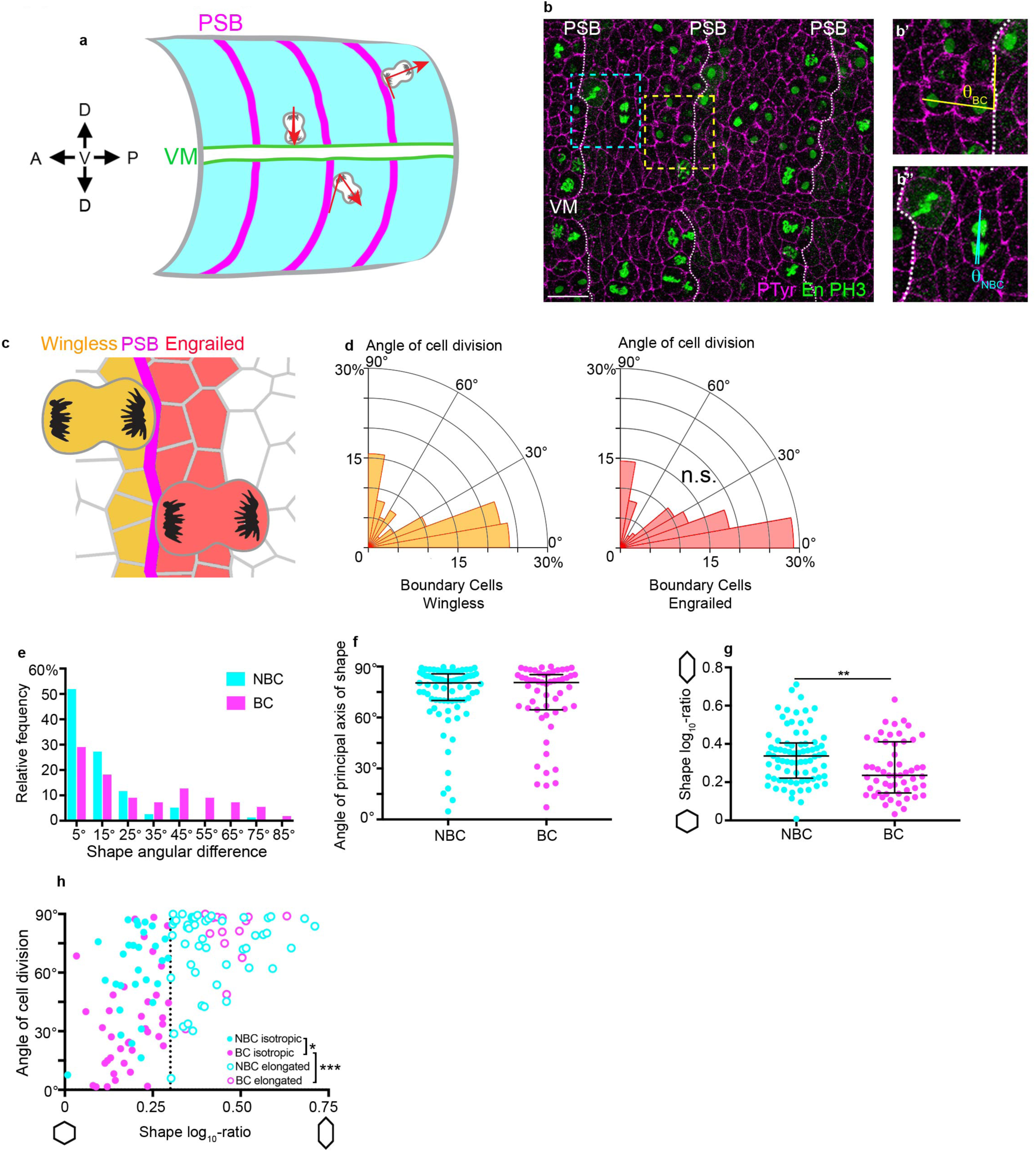
**a,** Diagram illustrating how the orientation of cell division is measured in fixed embryos. The angles given are relative to the orientation of the embryo’s antero-posterior axis. The angles are measured relative to the local curvature of the PSBs (which are DV-oriented interfaces) to take into account curvature artefacts due to embryo mounting. These angles are then converted by 90 degrees to obtain the orientation of cell division with respect to the antero-posterior axis. **b,** Example of an immunostaining used for angle measurements. A maximum projection of a confocal stack of a stage 9 embryo immunostained for phospho-Histone H3 (to highlight the chromosomes in dividing cells), Engrailed (to find the PSBs) and phospho-Tyrosine (to label the cell shapes) is shown. Note that the angle of cell division is measured in anaphase or telophase cells because by then the spindles have finished rotating (see Fig. 5). PSBs are highlighted by a dashed line. VM, ventral midline. Scale bar 20 µm. **b’**, close up of a BC (at telophase). **b’’**, close-up of a NBC (at anaphase). **c-d,** The orientation of dividing boundary cells was analysed according to their position relative to the PSB (either wingless or engrailed-expressing boundary cells). No significant difference was found between the two populations (Wg, *n*=165; En, *n*=124; Mann-Whitney test, *U*=9569, P=0.347). **e,** histogram of the angular differences between the orientation of cell division and the orientation of interphase cell shape for NBC (blue) and BC (pink) (see corresponding cumulative histogram in Fig. 1i). **f,** The principal axis of interphase cell shape is oriented perpendicular to the antero-posterior axis of the embryo for both NBC and BC populations (NBC, *n*=77; BC, *n*=55; Mann-Whitney test, *U*=2007, P=0.613). Median ± interquartile range shown. **g**, BC have less elongated shapes than NBC (NBC, *n*=77; BC, *n*=55; Mann-Whitney test, *U*=1549, P=0.008). Median ± interquartile range shown. **h** Cell division orientation as a function of cell elongation (cell shape log_10_-ratio). Above a log_10_-ratio threshold of 0.3 (DV/AP aspect ratio of 2), both NBC and BC behave similarly (NBC, *n*=48; BC, *n*=16; Kruskal-Wallis test, *H*=45.65, P=0.6393), dividing along DV. Below 0.3 however, NBC and BC behaviours are significantly different (NBC, *n*=29; BC, *n*=39; Kruskal-Wallis test, *H*=45.65, P=0.0194). For BC, the angle of cell division between elongated and isotropic cells was significantly different (Elongated BC, *n*=16; Isotropic BC, *n*=39; Kruskal-Wallis test, *H*=45.65, P<0.0001).

**Supplementary Figure 2:**
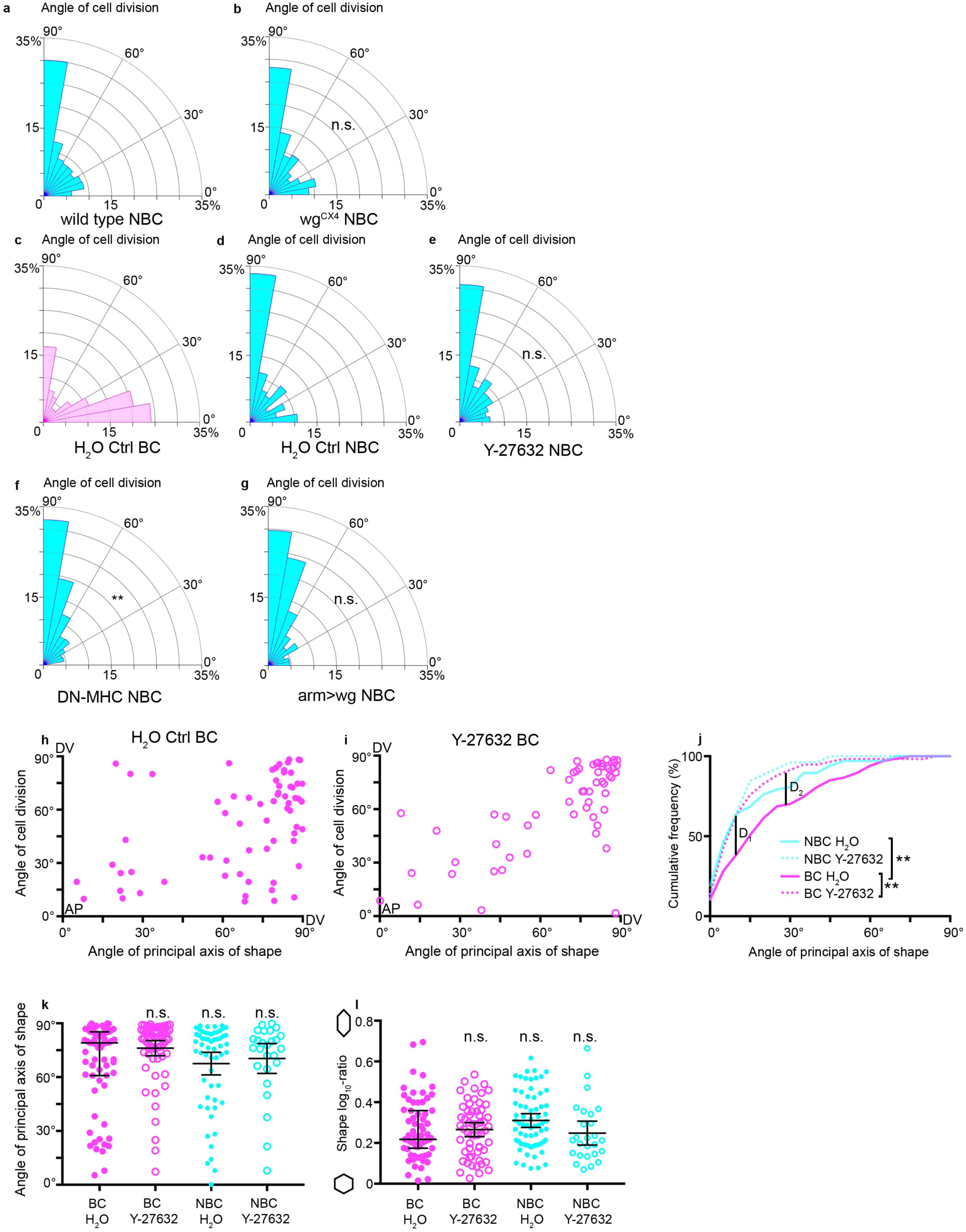
Histograms of NBC division orientation for the following genotypes: a, wild type (*n=*589) **b,** wg^CX4^ (*n=*618) for stages 9 to 11 embryos (Mann-Whitney test, *U*=128400, P=0.272). **c-e,** Histograms of BC (c, *n*=83) or NBC (d, *n*=132) division orientation in embryos injected with H_2_O or the ROK inhibitor Y-27632 (e, *n*=117) (Kruskal-Wallis tests, H_2_O BC vs H_2_O NBC, *H*=24.28, P<0.0001; H_2_O NBC vs Y-27632 NBC, *H*=24.28, P=1.0). **f,** Histograms of NBC division orientation for embryos expressing DN-MHC (*n*=454; wild type *n=*391 (from Fig. 1); Mann-Whitney test, *U*=78689, P=0.0044). **g,** Histograms of NBC division orientation for embryos expressing *arm>wg* (*n=*212; *n*=391, wild type (from Fig. 1); Kruskal-Wallis test, *H*=211.2, P=0.091). **h,i** BC from embryos injected with Y-27632 follow the long axis rule displaying a linear correlation between the angle of cell shape and the angle of cell division orientation (*n*=58; Spearman’s rho test, *r*=0.62, P<0.001) better than BC from H_2_O injected embryos (*n=*67; Spearman’s rho test, *r*=0.42, P<0.001). **j,** Cumulative histogram of the cell shape angular difference for BC H_2_O (*n=*67), BC Y27632 (*n=*58), NBC H_2_O (*n=*67), NBC Y-27632 (*n=*26). H_2_O BC are significantly different from H_2_O NBC (Kolmogorov-Smirnov test, *D*_*1*_=0.29, P=0.0051). H_2_O BC significantly different from Y-27632 BC, with a higher proportion of cells dividing according to their interphase shape (Kolmogorov-Smirnov test, *D*_*2*_=0.32, P=0.0031). All other comparisons are not significantly different. **k,** Shape principal axis orientation for H_2_O or Y-27632 treated BC and NBC (BC H_2_O, *n=*67; BC Y27632, *n=*58; NBC H_2_O, *n=*67; NBC Y27632, *n=*26). Kruskal-Wallis tests, all comparisons are not significantly different. Median ± interquartile range shown. **l,** Shape log_10_-ratio for H_2_O or Y-27632 treated BC and NBC (BC H_2_O, *n=*67; BC Y27632, *n=*58; NBC H_2_O, *n=*67; NBC Y27632, *n=*26). Kruskal-Wallis tests, all comparisons are not significantly different. Median ± interquartile range shown.

**Supplementary Figure 3:**
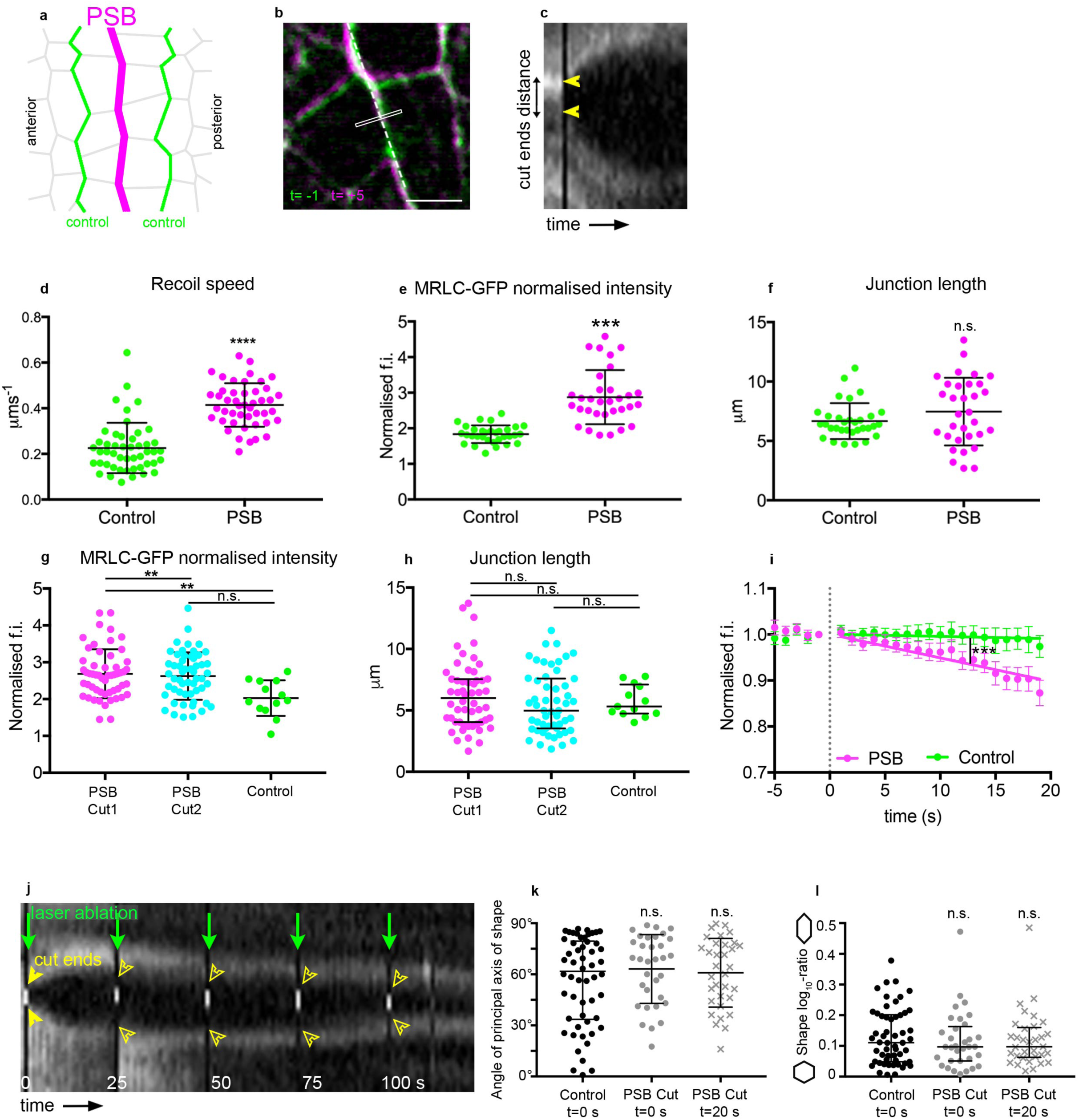
**a-d** Tension is higher at PSB compared to non boundary interfaces of boundary cells. (a) Tension at the PSB(magenta) is compared to that of adjacent control non boundary interfaces (green) in MRLC-GFP expressing embryos. (b) Overlay of a PSB before and after laser ablation (white rectangle, cut site). Scale bar 5 µm. (c) Kymograph spanning the dashed line in (b) used to measure the distance between cut ends over time (arrowheads). (d) Speed of recoil upon ablation (PSB, *n*=48; Control, n=46; Mann-Whitney test, *U*=209.5, P<0.0001). Mean ± SD shown. **e** Normalised fluorescence intensity for ablated control and PSB interfaces (PSB, *n*=48; Control, *n*=46; Student’s t-test, *t*=7.35, p<0.0001). Mean ± SD shown. **f** Junction length of ablated control and PSB interfaces (PSB, *n*=48; Control, *n*=46; Mann-Whitney test, *U*=442, P=0.351). Mean ± SD shown. **g** Normalised fluorescence intensity for ablated control and PSB interfaces at t=0 for consecutive PSB cuts (PSB Cut1, *n*=52; PSB Cut2, *n*=52; Control, *n*=29; One-way Anova, *F*=5.743, P=0.0042; Dunnett’s test for PSB Cut1 vs Control, P=0.0022; for PSB Cut2 vs Control, P=0.0059). Means ± SDs shown. **h** Junction length of ablated control and PSB interfaces for consecutive PSB cuts (PSB Cut1, *n*=52; PSB Cut2, *n*=52; Control, n=29; Kruskal-Wallis tests; PSB Cut1 vs Control, *H*=1.68, P>0.9999; PSB Cut2 vs Control *H*=1.68, P=0.922). Median± interquartile range shown. **i** Myosin intensity is decreased at the actomyosin cable after PSB ablation, compared to control interfaces anterior or posterior to the ablated region. Measurements are normalised to the initial fluorescence intensity before ablation. Means ± s.e.m.s shown. Slopes are significantly different (ANCOVA on linear regression, *F*=130, P<0.0001). **j** A representative kymograph for a PSB-cut experiment. Laser ablation was repeated every 25 s to prevent wound healing (green arrows). Lack of recoil upon repeated cuts was used to estimate successful loss of tension at the PSB (compare curve highlighted by the full yellow arrowhead, which shows recoil of cut ends, with the hollow yellow arrowheads, which show lack of recoil). **k** Orientation of the cell shape principal axis for control and PSB-cut treated cells at t=0s or at t=20s after ablation (Control, *n*=54; PSB-cuts, *n*=33; Kruskal-Wallis tests, *H*=1.268, all pair-wise comparisons P>0.50). Medians ± interquartile ranges shown. **l** Cell shape log_10_-ratios for control and PSB-cut treated cells at t=0s or at t=20s after ablation (Control, *n*=54; PSB-cuts, *n*=33; Kruskal-Wallis test, *H*=0.6968, all pair-wise comparisons P>0.99). Medians ± interquartile ranges shown.

**Supplementary Figure 4:**
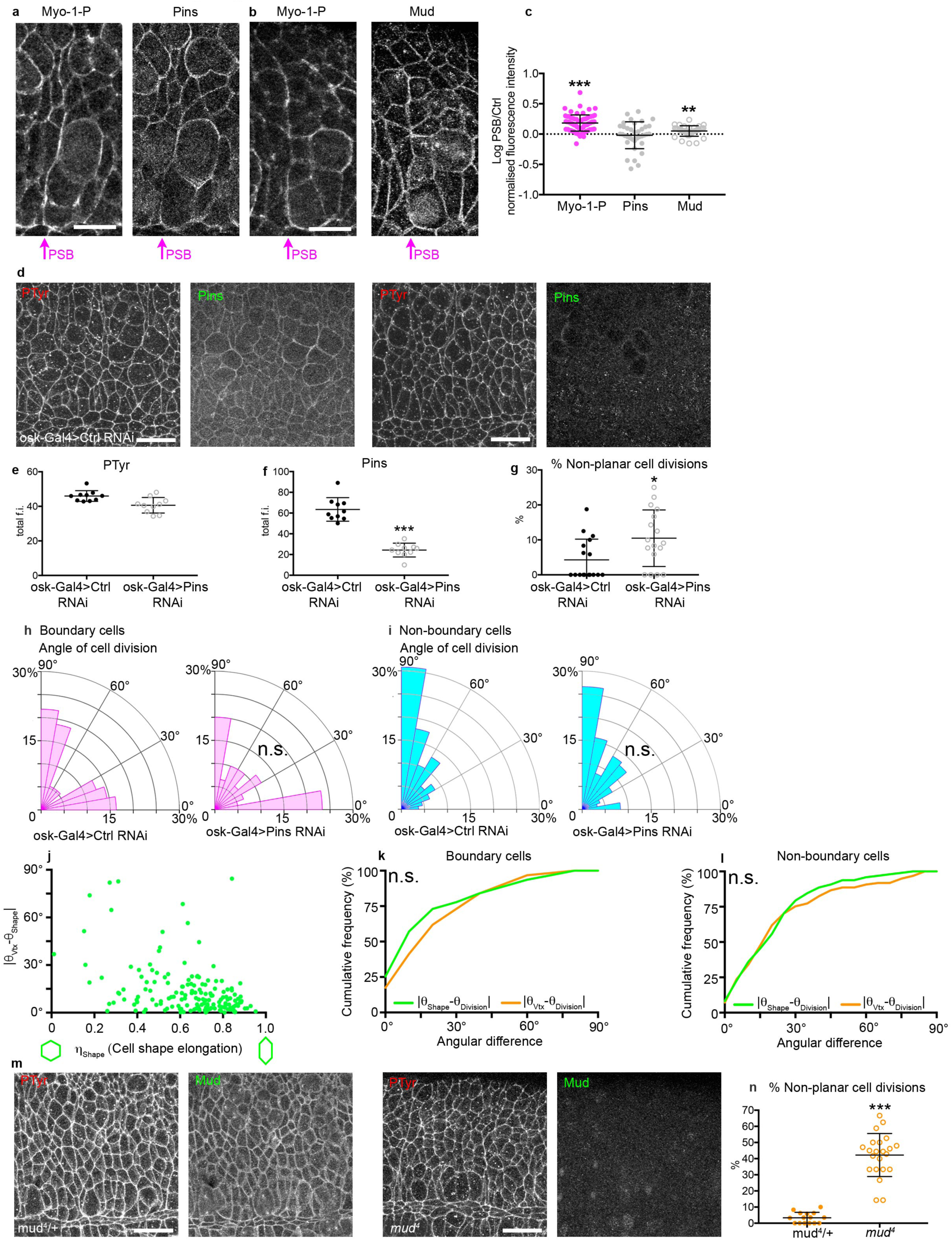
**a-c**, Endogenous Myosin-1-P and Pins localization in wild type embryos (a). Endogenous Myosin-1-P and Mud localization in wild type embryos (b). (c) Quantitation of Myosin-1-P, Mud and Pins fluorescence intensity enrichment at boundary interfaces compared to non-boundary interfaces (*n=*71, Myo-1-P interfaces; *n=*37, Pins; *n=*34, Mud; Wilcoxon signed-rank tests; Myo-1-P, P<0.0001; Pins, P=0.988; Mud, P=0.0019). Means ± SDs shown. **d-g** Endogenous Pins and PTyr immunostainings for Control RNAi and Pins RNAi embryos. Scale bar 20 µm (d). Quantitation of PTyr (e) and Pins (f) total fluorescence intensity (*n*=10 embryos, Ctrl RNAi; *n*=10 embryos, Pins RNAi; unpaired Student’s *t*-tests; PTyr, *t*=3.09, P=0.0064; Pins, *t*=9.40, P<0.0001). Means ± SDs shown. **g** Percentage of cells dividing out of the epithelial plane for Control (9/201 cell divisions) or Pins (21/177 cell divisions) RNAi (*n*=17 embryos per genotype, Mann-Whitney test, *U*=80.5, P=0.0216). Means ± SDs shown. **h**, Histogram of cell division orientation for boundary cells (BC) in Pins RNAi (*n*=60), or Control RNAi (*n*=55; Mann-Whitney test, *U*=1568, P=0.649) embryos. **i,** Histogram of cell division orientation for non-boundary cells in Pins RNAi (*n*=107), or Control RNAi (*n*=120; Mann-Whitney test, *U*=5881, P=0.276) embryos. **j** Scatterplot of the absolute difference between cell shape principal axis orientation and vertex cluster orientation, |θ_TCJ_-θ_Shape_|, against cell elongation (η_Shape_). Note that less elongated cells have a higher angular difference between shape and vertex orientation. **k-l,** Cumulative histograms of the Shape deviation angle or Vertex deviation angle for BC (a, *n=*63 from 3 embryos; Kolmogorov-Smirnov test, *D*=0.1579, P=0.972) and NBC (b, *n=*97 from 3 embryos; Kolmogorov-Smirnov test, *D*=0.0824, P=0.896). **m,** Endogenous Mud and PTyr immunostainings for mud^4^/+ and *mud^4^* embryos. Scale bar 20 µm. **n,** Percentage of cells dividing out of the epithelial plane for mud^4^/+ (12/277 cell divisions) or *mud^4^* (124/289 cell divisions) embryos (*n*=15 embryos, *mud^4^*; *n*=22 embryos, *mud^4^/*+; Student’s *t*-test, *t*=11.01, P<0.0001). Means ± SDs shown.

**Supplementary Figure 5:**
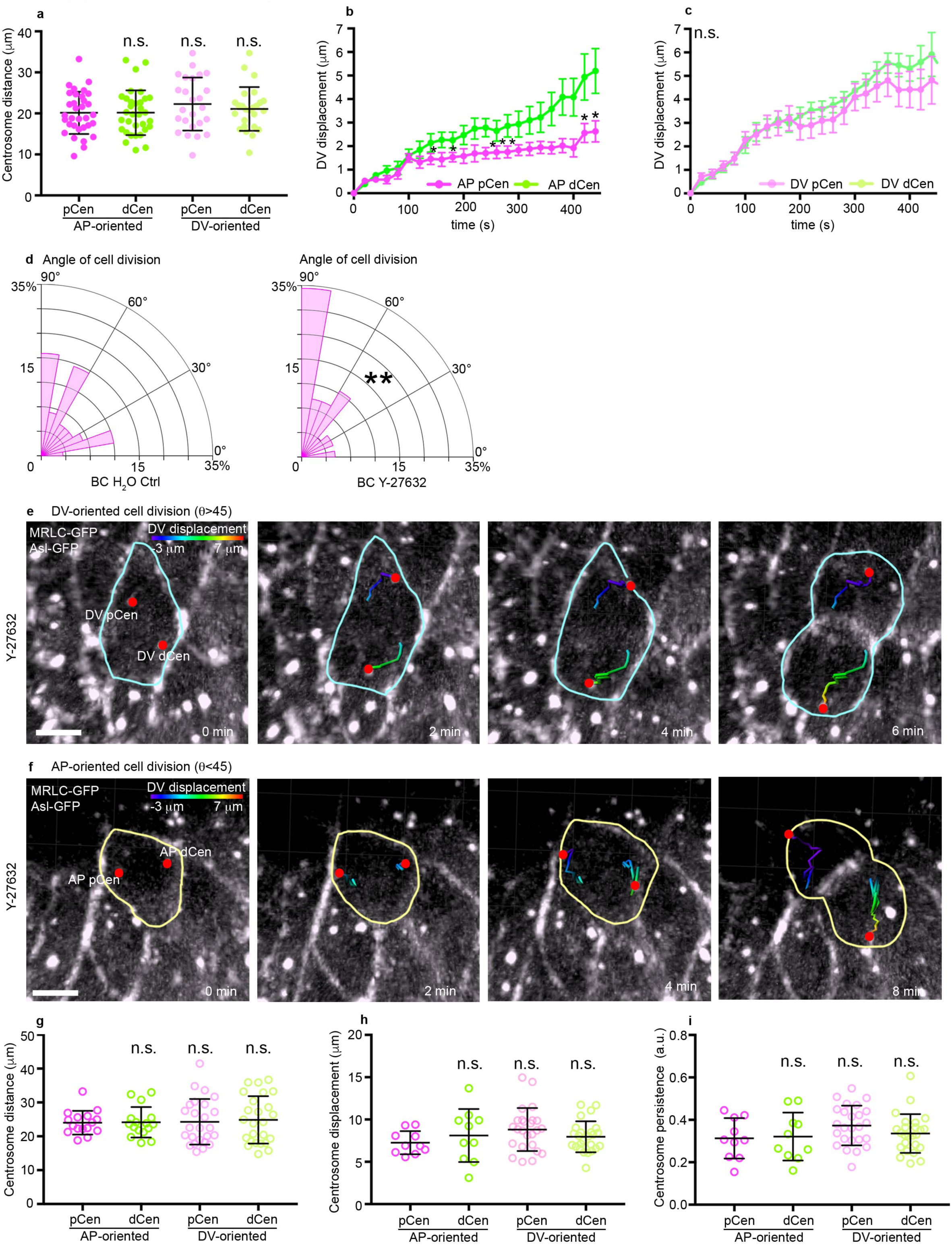
**a** Total distance travelled by each centrosome from NEBD to cytokinesis (*n=*33, AP pCen; *n=*33, AP dCen; *n*=25, DV pCen; *n*=25, DV dCen; Kruskal Wallis tests, *H*=2.904, all pair-wise comparisons P>0.75). Means ± SDs shown. **b** Absolute displacement along the DV axis of the embryo over time (t=0 NEBD) for AP-Oriented BC divisions (*n=*16, AP pCen; *n=*16, AP dCen; paired Student’s t-tests, P<0.05 shown by asterisk). Means ± s.e.m.s shown. **c** Absolute displacement along the DV axis of the embryo over time (t=0 NEBD) for DV-Oriented BC divisions (*n=*19, DV pCen; *n=*19, DV dCen; paired Student’s t-tests, P>0.05). Means ± s.e.m.s shown. **d** Histogram of OCD for BC from embryos expressing MRLC-GFP and Asl-GFP injected with either H_2_O or Y-27632 (*n=*67, BC H_2_O; *n=*58, BC Y-27632; Mann-Whitney test, *U*=1525, P=0.0384). **e,f** Representative DV-oriented (e) and AP-oriented (f) BC cell division from an Y-27632 injected embryo expressing MRLC-GFP and Asl-GFP. Centrosome tracks are highlighted and colour-coded for DV displacement. Scale bar 5 µm. **g** Total distance travelled by each centrosome from NEBD to cytokinesis for Y-27632 injected embryos (*n=*10, AP pCen; *n=*10, AP dCen; *n*=25, DV pCen; *n*=25, DV dCen; One-way Anova, F=0.0813, P=0.97, all pair-wise comparisons P>0.90). Means ± SDs shown. **h** Total displacement for each centrosome from NEBD to cytokinesis for Y-27632 injected embryos. (*n=*10, AP pCen; *n=*10, AP dCen; *n*=25, DV pCen; *n*=25, DV dCen; One-way Anova, *F*=1.278, P=0.28, all pair-wise comparisons P>0.15). Means ± SDs shown. **i** Persistence for each centrosome from NEBD to cytokinesis for Y-27632 injected embryos (*n=*10, AP pCen; *n=*10, AP dCen; *n*=25, DV pCen; *n*=25, DV dCen; Kruskal Wallis tests, all pair-wise comparisons P>0.30). Means ± SDs shown.

**Supplementary Figure 6:**
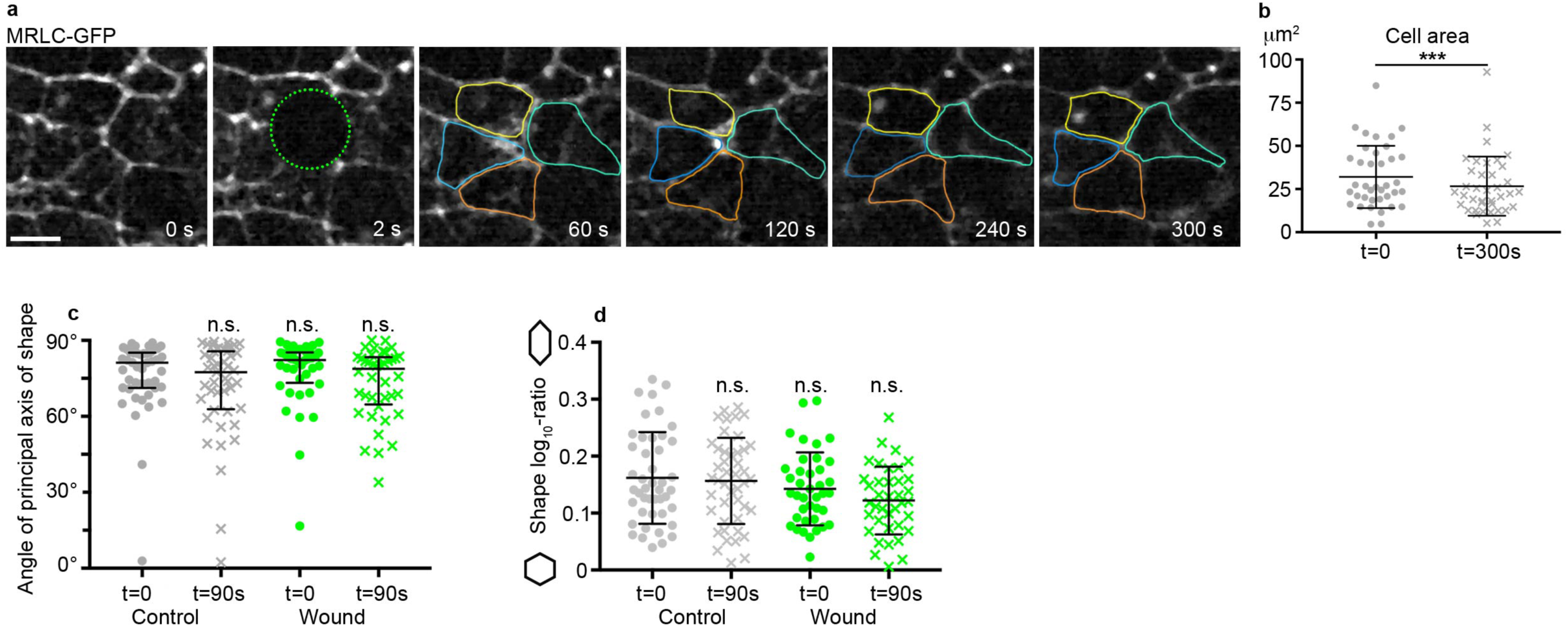
**a-b** IR laser irradiation leads to wound healing without triggering cell delamination. (a) Still images from a gain of tension experiment. Green circle: site of wounding. Cell outlines are highlighted, scale bar 5 µm. (b) Cell area at t=0 and at t=300s after wounding (*n*=38; Wilcoxon matched pairs signed-rank test, P=0.001). Means ± SDs shown. **c** Orientation of the cell shape principal axis for control and wound treated cells at t=0 and at t=90 seconds after wounding (*n*=45 Sham, *n*=40 Wound; Kruskal-wallis tests, all pair-wise comparisons P=1.0). Means ± interquartile ranges shown. **d** Shape log_10_-ratio for control and wound treated cells at t=0 and at t=90s after wounding (*n*=45, control; *n*=40, wound; One-way Anova, *F*=2.52, P=0.060; Sidak’s multiple comparisons, all pair-wise comparisons P>0.05). Means ±SD shown.

## METHODS

### Fly Stocks

The following stocks were used in this study: yw^67^ (BDSC, #6599) was used as wild type. The stock sqh^AX3^; sqh-sqhGFP42;GAP43^mem^::mCherry/TM6B^1^ was used for laser ablation and wound healing experiments. The stock sqh^AX3^; shotgun::GFP^2^, sqh::mCherry^3^ was used for live imaging and automatic tracking of cell divisions. The following transgenic lines were used: en-lacZ^4^ ; arm-Gal4^5^; UAS-wg^6^; arm-FRT-stop-FRT-Gal4::VP16^5^; KB19 (male-specific flipase)^5^; UAS-GFP::DN-MHC^7^; ubi-E-Cadherin-GFP^8^; Jupiter::mCherry^9^; en-Venus^10^, osk-Gal4::VP16 (BDSC, #44242); pins RNAi (BDSC, #37479); eGFP RNAi (BDSC #41556); Asl-Asl::GFP^11^; *GFP-Mud62E1.GFP-Mud65B^12^*. The following null mutant alleles were used: wg^cx4^ ^13^, mud^4^ ^14, 15^, Asl^B46^ ^16^. Note that in the main text, Sqh (encoding *Drosophila* Myosin Regulatory Light Chain) is indicated throughout the text as MRLC and Shotgun (encoding *Drosophila* E-Cadherin) as E-Cadherin.

### Genotypes

<u>Figure 1</u>(b-e)-Figure S1(d) *yw*^67c23^. Figure 1(f-j)-Figure S1(e-g) *en-Venus/ ubi-E-Cadherin-GFP; Jupiter::-mCherry/+*. <u>Figure 2</u>(a, d) *sqh^AX3^;sqh-sqhGFP42;GAP43^mem^::mCherry/TM6B.* (e) *wg^cx4^,enLacZ* (left panel), *yw^67^*(right panel). (f) *yw^67^* (left panel), *arm-Gal4::VP16/UAS-DN-MHC::GFP* (right panel).(g) *arm-Gal4/UAS-wg*. The same genotypes were used, as indicated, in FigureS2 (a-b). <u>Figure S2</u> (c-g) *sqh^AX3^/+;sqh-sqhGFP42/Asl-Asl::GFP; GAP43^mem^::mCherry/Asl^B46^*. <u>Figure3-FigureS3</u> *sqh^AX3^;sqh-sqhGFP42;GAP43^mem^::mCherry/TM6B.* <u>Figure 4</u> (b-d)-Figure S4(a-b) *sqh^AX3^; shotgun::GFP; sqh::mCherry*. (e-e’)-Figure S4(c-f) *yw^67^*. (f) *GFP-Mud62E1.GFP-Mud65B2/TM6B.* (g-h)-FigureS4 (g-j) *osk-Gal4::VP16/UAS-eGFP^RNAi^* or *osk-Gal4::VP16/UAS-Pins^RNAi^* as indicated. (i-j)-FigureS4(k-l) *mud^4^/FTG* or *mud^4^* as indicated. <u>Figure 5</u>(a-c) *en-Venus/ ubi-DE-Cadherin-GFP; Jupiter::mCherry/+.*(d-l)-Figure S5(a-g) *sqh^AX3^/+;sqh-sqhGFP42/Asl-Asl::GFP; GAP43^mem^::mCherry/Asl^B46^*. <u>Figure 6</u>-Figure S6. *sqh^AX3^; sqh-sqhGFP42;GAP43^mem^::mCherry/TM6B.*

### Immunostainings

Embryos collected in a basket from plates containing agar-apple juice were washed in tap water and dechorionated using commercial bleach for 2 minutes, rinsed and dried. Embryos were then fixed at the interface of a 1:1 solution of 37% formaldehyde : 100% heptane for 7 minutes, followed by either manual devitellinisation in PBS 0.1% Triton-X-100 (PTX) or by 100% methanol devitellinisation. Methanol devitellinised embryos were re-hydrated by sequential washes with 75% methanol/PBS, 50% methanol/PBS, 25% methanol/PBS and PBS. They were then blocked in 1% BSA in PTX for 30 minutes and incubated overnight with primary antibodies. Embryos were washed three times in PTX for 5 minutes before secondary antibody incubation for 1 hour at room temperature. Finally, they were washed three more times in PTX and mounted in Vectashield (Vector laboratories) for imaging.

### Antibodies

The following antibodies were used: Rabbit Phospho-histone H3 (Cell Signalling #9701, 1:200), rabbit anti-Engrailed (Santa Cruz Biotechnology d-300; 1:200), goat anti-GFP-FITC (Abcam ab6556, 1:500), guinea pig anti-Sqh-1P (1:100, a gift from R.E. Ward IV), mouse anti-phospho-Tyrosine (Cell signaling #9411; 1:1000), mouse anti-Wingless (DSHB 4D4; 1:50); mouse anti-Dlg (DSHB 4F3; 1:500) Rabbit anti-Pins^17^ (1:1000, a gift from F. Matsuzaki), rabbit anti-Mud^17^ (1:200, a gift from F. Matsuzaki). Secondary antibodies conjugated to fluorescent dyes were obtained from Jackson ImmunoResearch Laboratories, Invitrogen and Life Technologies. Cell nuclei were stained using DAPI (Sigma-Aldrich).

### Confocal imaging

Embryos were individually mounted in a ventral orientation under a tape bridge on either side of the slide, so that they were sufficiently flattened. Imaging was either performed on a Nikon Eclipse TE2000 microscope incorporating a C1 Plus confocal system (Nikon) and images captured using Nikon EZ-C1 software; or on a Leica TCS SP8 confocal microscope and images captured using LAS X software (Leica). Optical z-stacks were acquired with a depth of 0.5-1 µm between successive optical z-slices. All embryos were imaged using a violet corrected 60x oil objective lens (NA of 1.4). The gain and offset were optimized for each immunostaining and maintained the same between control and mutant conditions.

### Analysis of orientation of cell division in fixed embryos

To analyse cell division orientation in fixed embryos, embryos were immunostained with Phospho-histone H3 antibodies to label mitotic chromosomes, antibodies against Wg or En to identify the PSB and antibodies against membrane marker (PTyr or Dlg) to determine cell shapes. Only cells in anaphase or telophase were analysed. For each cell division, the angle θ_PSB_ (see Fig S1a-b) between the local curvature of the PSB and the separating chromosomes was measured using the angle tool using the Fiji software (NIH Image). Angles were always measured as acute angles. To obtain the orientation of cell division in relation to the ventral midline, which serves as a proxy for the AP axis of the embryo, the angle θ_PSB_ was transformed as follows: θ_midline_*=*90°-θ_PSB_, under the assumption that the midline would be always perpendicular to the PSB. An example of the θ_PSB_ measurement for a BC and a NBC is given in Supplementary Figure S1.

### Live imaging

Dechorionated embryos were transferred into halocarbon oil (Voltalef PCTFE, Arkema), mounted on a stretched oxygen-permeable membrane with their ventral side facing up, and covered by a coverslip which was supported by a single coverslip bridge on either side of the membrane. Imaging was performed using a Nikon Eclipse E1000 microscope equipped with a spinning disk unit (Yokogawa CSU10), laser module with 491nm and 561nm excitation (Spectral Applied Research LMM2), and a C9100-13 EM-CCD camera (Hamamatsu). Image acquisition was carried out using the Volocity software (Perkin Elmer). A frame delay of 20s with 0.7μm Z-intervals was used to image mitotic spindles and centrosomes, while images were acquired every 30 s with 1μm Z-intervals for automatic tracking.

### Automated tracking and cell shape versus tricellular vertex analysis

Cells were segmented and tracked in three movies with membrane and myosin fluorescence (E-Cadherin-GFP; MRLC-mCherry) as previously (Blanchard et al., 2009; Tetley et al., 2016). Embryos were imaged from late stage 8 to stage 10 for 2 to 3 hours. The timepoint of the first cell division in the mesectoderm was used to synchronise movies. Only ventral neurectoderm cell divisions, which start at stage 9, were analysed. PSBs were identified by inspection of the MRLC-mCherry signal. Mother cells were identified at the frame before abscission, when a new junction separated the mother into two daughter cells (*n*=277). The orientation of cell division was measured as the orientation between daughter centroids at abscission (*θ*_division_). For each mother cell we looked 12 minutes back in time, corresponding to approximately the start of NEBD. At 12 mins prior to abscission, the orientation (*θ*_shape_) and strength (*η*_shape_, between 0 and 1) of cell elongation and the orientation (*θ*_Vtx_) and strength (*η*_Vtx_, between 0 and 1) of vertex clustering were calculated according to published methods (Bosveld et al., 2016, Nature). Finally, we calculated the absolute degrees difference between the orientation of cell division and the orientations of both cell elongation and vertex clustering and compared these distributions.

### Analysis of orientation of cell division and interphase shapes in live embryos

To analyse whether cell divisions followed the interphase principal axis of cell shape, we imaged *ubi-DE-Cadherin/En>Venus; JupiterCherry/+* embryos. Embryos were imaged on a spinning disk microscope using a 100x magnification to better visualise the mitotic spindle. 21 movies (n=3 movies from stage 9 embryos, n=15 movies from stage 10 embryos, n=3 movies from stage 11 embryos) were analysed. The PSBs were identified by the fluorescent boundary marker En>Venus. To analyse whether cell divisions followed their principal axis of cell shape, interphase cell shapes were manually traced in Fiji at t=-12 minutes from the end of cytokinesis, corresponding roughly to the start of NEBD as described in the paragraph above. Cell principal axis orientation and cell shape longest and shortest axis were extracted by analysing the cell shape using the best-fit ellipse tool in Fiji. Cell shape orientation was measured with respect to the ventral midline of the embryo, similarly to cell division orientation, as shown in Fig. S1A. Cell shape log_10_-ratio was defined as the log_10_ of the ratio between the ellipse longest and shortest axis. Cell division orientation was measured at anaphase by tracing the angle between the mitotic spindle (for cells expressing Jupiter-mCherry, Fig. 1) or the two centrosomes (for cells expressing Asl-GFP, Fig. S2) and the local PSB curvature, as depicted in Supplementary Fig. 1a. To analyse the rotation of the mitotic spindle, the angle between the spindle and the local PSB curvature was measured every 20 s from NEBD to cytokinesis. NEBD was identified by inspecting the Jupiter-Cherry signal, since before NEBD Jupiter is excluded from the nucleus. The angular difference was defined as the |Shape Principal Axis Orientation-Cell Division Orientation|.

### Y-27632 Rho kinase inhibitor injections

Stage 8 SqhAX3/+; Asl-GFP/SqhGFP42; AslB46^16^/GAP43-mCherry embryos were mounted with their ventral side facing a glass coverslip with heptane glue, covered with halocarbon oil and injected through the posterior into the yolk at room temperature with 1 mM Y27632 (TOCRIS) in dH_2_O, or dH_2_O in control experiments^18^. This concentration of Y-27632 does not affect epithelial integrity neither it does inhibit cytokinesis^18^. Embryos were allowed to recover for 30 minutes at 18**°**C, then imaged for ∼2 hours at 21**°**C.

### Laser ablation

Laser ablation experiments were performed using a TriM Scope II Upright 2-photon Scanning Fluorescence Microscope controlled by Imspector Pro software (LaVision Biotec) equipped with a tuneable near-infrared (NIR) laser source delivering 120 femtosecond pulses with a repetition rate of 80 MHz (Insight DeepSee, Spectra-Physics). The laser was set to 927nm, with power ranging between 1.40-1.70 W. The maximum laser power reaching the sample was set to 220 mW and an Electro-Optical Modulator (EOM) was used to allow microsecond switching between imaging and treatment laser powers. Laser light was focused by a 25x, 1.05 Numerical Aperture (NA) water immersion objective lens with a 2mm working distance (XLPLN25XWMP2, Olympus). Ablations were carried out during image acquisition (with a dwell time of 9.27 µsec per pixel), with the laser power switching between treatment and imaging powers as the laser scanned across the sample. Targeted line ablations of ∼2 µm length were performed at the centre of junctions on the PS boundary or on control, non boundary dorso-ventral (DV) oriented junctions, using a treatment power of 220 mW. Images were acquired with a frame delay of 731 ms, more than 45 ablations per condition were carried out, 2-4 ablations per embryo. For consecutive ablations, line ablations of ∼2 µm length were performed at the centre of junctions on a PS boundary, and after 20 s a second ablation was carried out on the same PSB at a distance of two junctions (one cell, see Fig. 3e) from the first cut, as previously described^19^. Images were acquired with a frame delay of 1s, more than 45 ablations per condition were carried out, 2-4 ablations per embryo. For loss of tension experiments, line ablations of ∼2 µm length were performed at the centre of junctions on a PS boundary next to a dividing cell in metaphase. Ablations were repeated every 24 seconds to prevent tissue healing, and after imaging kymographs were inspected to verify loss of recoil upon consecutive ablations (arrows, Fig. S3f). Control ablations were carried out by setting the EOM unit treatment power at 25% of 220 mW (the same intensity used to image the sample) instead of 100%. As for treatment ablations, control ablations were also repeated. Images were acquired with a frame delay of 1s.

### Laser ablation analysis

To analyse recoil velocities, images were background subtracted and denoised using Fiji. A kymograph spanning the ablated region was generated using the dynamic reslice function in Fiji, and the distance between the two ends of the cut was measured up to 20 seconds after ablation. Linear regression was performed on the first 5 timepoints after ablation using a custom-made Matlab script^20^ and the slope of the regressed line was used^1^ a measure of the cut ends recoil velocity. Junction length for PSB and non-PSB interfaces was measured using Fiji, and Myosin intensity was quantified in Fiji by drawing a 3 pixel-wide line selection on the junction of interest at t=0 and normalising it by dividing it by the signal of Myosin in the cytoplasm of the same cell at the same timepoint.

To measure Myosin signal intensity over time (Fig. S3e, Fig 6b), a 3 pixel-wide line selection on the junction of interest or a ∼5µm diameter circle on the wounded/control area was drawn in Fiji and its intensity was measured and normalised by subtracting the mean grey value of the whole imaged area for each timepoint to correct for sample bleaching.

### IR-laser wounding

Circular ablations of ∼5µm diameter were performed twice with a 1 second interval on non boundary junctions next to a metaphase non-boundary cell, using a EOM treatment power of 80% of 220 mW. To measure tension upon wound healing, a line ablations of ∼2 µm length were performed at the centre the wounded area 90 seconds after the circular ablation, which corresponds to the peak of Myosin intensity (see Fig.6b) and the recoil speed was measured as described. Images were acquired with a frame delay of 2s.

### Quantifications from immunostainings

Quantifications were carried out on maximum intensity projections, which were derived from the minimum number of z-slices needed to contain all the signal. To quantify whether Pins or Mud were enriched at the PSB, the position of the PSB was identified by counterstaining with anti-En or anti-Wg antibodies. PSB or non-PSB interfaces were traced as 3-pixel wide lines and the fluorescence intensity of the selection was normalized to the modal grey value of the embryo as previously described^21^. To quantify the extent of Pins knockdown, control RNAi and Pins RNAi images were acquired in the same session using the same confocal settings. Maximum intensity projections were generated for each channel, the outline of the embryo was drawn and the absolute fluorescence intensity of the control channel (P-Tyrosine) or the experiment channel (Pins) was measured and plotted as previously described^21^.

### Centrosome tracking

Movies from H_2_O or Y27632-injected SqhAX3/+; Asl-GFP/SqhGFP42; AslB46/GAP43-mCherry embryos were analysed using the Imaris software (Bitplane). First, movies were corrected for rotational and translational xyz drift. Boundary cells were identified by inspection of the MRLC-GFP signal and individual centrosomes were manually tracked along the 3 xyz dimensions from NEBD to cytokinesis. NEBD was identified by inspecting the MRLC-GFP signal, since before NEBD Myosin is partially excluded from the nucleus^22^. Measurements such as centrosome speed, distance, displacement and persistence were calculated in Imaris, exported and analysed using Excel (Microsoft) or GraphPad Prism.

### Statistical analysis

Statistical analysis was performed using GraphPad Prism. Angular histograms were plotted using a custom-made R script. Data from quantifications are reported as mean±SD, mean±s.e.m., median±25^th^/75^th^ percentiles or histograms according on whether they follow a normal distribution or not. On normally distributed data, two-tailed Student’s t-tests (two experimental groups) or One-way Anova followed by Dunnett’s multiple comparisons test (multiple experimental groups) were performed, while for non-normally distributed datasets Mann-Whitney, Kolmogorov-Smirnov (two experimental groups) or Kruskal-Wallis multiple comparisons tests (multiple experimental groups) were performed as described in the figure legends. Statistical analysis of cell division orientation histograms was carried out using two-tailed Mann-Whitney non-parametric tests^23, 24^. For all tests, a confidence level of 0.05 was considered statistically significant.

